# Single-molecule observation of multi-scale dynamic heterogeneity in the molecular bearing of the bacterial flagellum

**DOI:** 10.1101/2025.02.20.639300

**Authors:** Martin Rieu, Ashley L Nord, Alexis Courbet, Hafez El Sayyed, Richard M Berry

**Affiliations:** Department of Physics, University of Oxford, Oxford UK; Kavli Institute for Institute for Nanoscience Discovery, Oxford UK; Centre de Biologie Structurale, Université de Montpellier, CNRS, INSERM. Montpellier, France; Institute for Protein Design, University of Washington, Seattle, WA, 98105, USA; Department of Biochemistry, University of Washington, Seattle, WA, 98195, USA; Howard Hughes Medical Institute, University of Washington, Seattle, WA, 98105, USA

## Abstract

The bacterial flagellar motor (BFM) is a protein-based rotary machine that drives bacterial motility. It comprises a stator anchored to the cell wall and a rotor in the cytoplasmic membrane, linked via the flagellar rod to the extracellular hook and filament. We describe high-speed polarization microscopy of gold nanorods attached to the hook, allowing unprecedented continuous recording of single BFMs with ∼0.5° resolution at 250 kHz for tens of minutes. The angular distribution of actively rotating motors exhibited multiple periodicities that are consistent with structure of the bearing between the rod and LP-ring. We observed passive diffusional rotation of this bearing, revealing complex dynamics at odds with the expected classical Ornstein-Uhlenbeck process in a smooth potential. Resolved transitions between the main 26-fold bearing states exhibited highly non-Poissonian kinetics spanning four orders of magnitude in timescale. Using the average transition times, we estimated the interaction barrier in the bearing to be close to 13 k_B_T, suggesting that the previously estimated torque generated by a single stator unit is just sufficient to overcome this barrier. At sub-millisecond timescales we observed anomalous ultra-slow diffusion typically associated with multi-scale disordered systems, despite the ordered crystalline atomic structure of the bearing revealed by cryo-Electron Microscopy. Over longer periods, we observed dynamic shifts in the preferred angular positions, indicating that the motor’s potential energy landscape evolves over time. We speculate that these newly discovered rich dynamics may be a universal feature of protein machines.

## Introduction

The fundamental processes of life are performed by dynamic biological nano-machines, driven and dominated by thermal fluctuations at the molecular scale^1^. At this level solid, smooth surfaces do not exist. The molecular machines of life are made of discrete, lumpy protein units interacting at the atomic level through Van der Waals and hydrophobic forces, electrostatic potentials and hydrogen bonds. Recent advances in cryo-electron microscopy are revealing intricate atomic structures of ever larger and more complicated molecular machines, leading in turn to ever more detailed models of how these machines work^2–10^. But the structures are obtained by averaging ensembles of snapshots of frozen, isolated complexes - leaving dynamics to be guessed qualitatively or occasionally inferred from differences between alternative structures of the same machine^11^. The theory of thermally driven dynamics dates back to foundational contributions by Arrhenius (1889), Eyring (1935), and Kramers (1940). Arrhenius’ law (Arrhenius 1889) empirically describes the dependence of reaction rates on temperature in terms of an activation energy barrier. Eyring’s transition state theory^12^ provides a molecular-level explanation, describing rare, thermally activated jumps between discrete states separated by energy barriers. Kramers’ theory^13^ extends these ideas by modeling systems as freely diffusing along smooth, one-dimensional energy landscapes defined by a reaction coordinate, allowing transition rates to be calculated based on local curvature and friction. Since then, theorists and experimentalists^14^ have investigated how the shape of the energy landscape can alter transition rates for a given barrier height and length, showing that intermediate states and rough landscapes could accelerate transitions; with applications to the understanding of protein folding^15,16^, protein assembly^17^, or to the detection of intermediate states from the long tail of the distribution of transition times^18^. However, experimental methods with sufficient resolution and lifetime to characterize the dynamics of a large biomolecular machine have been lacking.

Here we used single-molecule measurements with unprecedented angular resolution to reveal rich dynamics of the molecular bearing of the bacterial flagellum, over timescales ranging from ∼10 μs to ∼10 minutes. The flagellum contains one of the best-studied of all large molecular machines: an ion-driven rotary motor ∼50 nm in diameter embedded in the bacterial envelope, containing hundreds of proteins of tens of different types, called the Bacterial Flagellar Motor (BFM)^4,19,20^. Recent advances in cryo-electron microscopy have revealed atomic structures of several parts of the BFM^5,6,8,21–24^ including the bearing that transmits rotation of the C-ring in the cytoplasmic membrane through the cell wall and outer membrane to the extracellular hook and filament in *Salmonella Enterica typhimurium*^5,25^. The bearing structure shows the 26-fold LP-ring tightly surrounding the flagellar rod, a polymer built on a helical lattice with 1-, 5-, 6- and 11-start helices. We directly observed the one-dimensional reaction coordinate of the bearing, the rotation angle between the LP-ring and the rotating flagellar rod. Rods were labelled by individual gold nanorods attached via the flagellar hook, and we calculated their orientation from the polarization anisotropy of laser illumination backscattered by nanorod surface plasmon resonance^26–28^ sampled at 250 kHz in a custom-built darkfield microscope.

We saw the undriven rotation of the bearing dominated by transitions between energy minima typically separated by approximately 1/26^th^ of a revolution, consistent with the LP-ring symmetry and previous observations of stepping in powered BFMs^5,25,29^. Between transitions we observed ultra-slow anomalous diffusion, a signature of rough local energy landscapes. Transition times approximated a power-law distribution over a very large range of timescales, from a few hundred microseconds to a few seconds. Furthermore, we resolved both dynamic and static heterogeneity of the bearing: the transition rates between energy minima varied between different bearings, between different locations in the same bearing and even over time at the same location in a single bearing. Also, the locations and spacing of energy minima varied both between bearings and over time in a single bearing. This unexpected rich dynamic behaviour adds a totally new level to our understanding of this canonical large molecular machine, complementary to the static atomic precision of the averaged cryo-EM structure.

We also measured the symmetries of the flagellar bearing. The distribution of dwell angles showed a strong ∼26-fold component and varied between individual bearings, consistent with our observations of passive bearing rotation. We computed the bearing potential as a function of rotation angle using Rosetta^30^ and the cryo-EM structure^31^, which indicated a rough landscape with variable 26-fold barriers. We also used a simple simulation to predict how (i) the observed dynamic and static heterogeneities and (ii) other periodicities in the dwell distributions might arise from (i) the tight molecular bearing and (ii) symmetry mismatches, observed in the atomic structure of the bearing.

## 2 Results

### 2.1 High resolution measurement of flagellar rotation using polarization anisotropy of gold nanorods

To perform fast measurements of flagellar rotation we took inspiration from experiments in which the dipolar polarization pattern of an asymmetric emitter encodes its orientation^27,32–35^ in 4 polarization channels. This compact encoding allows much faster acquisition rates over much longer durations than data-intensive video methods. The method is also insensitive to noise arising from the position of the emitter, and unlike single fluorescent molecules, nanorods do not photobleach, allowing indefinite observation of each nanorod. We attached gold nanorods to the biotinylated hook^36^ of the BFM (Figure 1A) and custom-built a laser backscattering dark-field microscope to measure their orientation (Figure 1B). Circularly polarized collimated laser illumination reaches the sample through a hole in a mirror, which also lets light reflected by the coverslip escape back towards the laser. Light back-scattered by the rod is reflected to a combination of polarizing and non-polarizing beamsplitters which separate the signal into four polarizations (0°, 45°, 90°, 135° from vertical; Figure 1C) and direct them onto four avalanche photo diodes. Measuring 45° and 135° polarizations is necessary to remove the degeneracies of the signal close to the 0° and 90° axes. A single Berek Compensator reverses phase shifts between 0° and 90° components, accumulated from reflections at dielectric mirrors in the light-path, which would otherwise mix the intensities of the 45° and 135° components.

**Figure 1:**
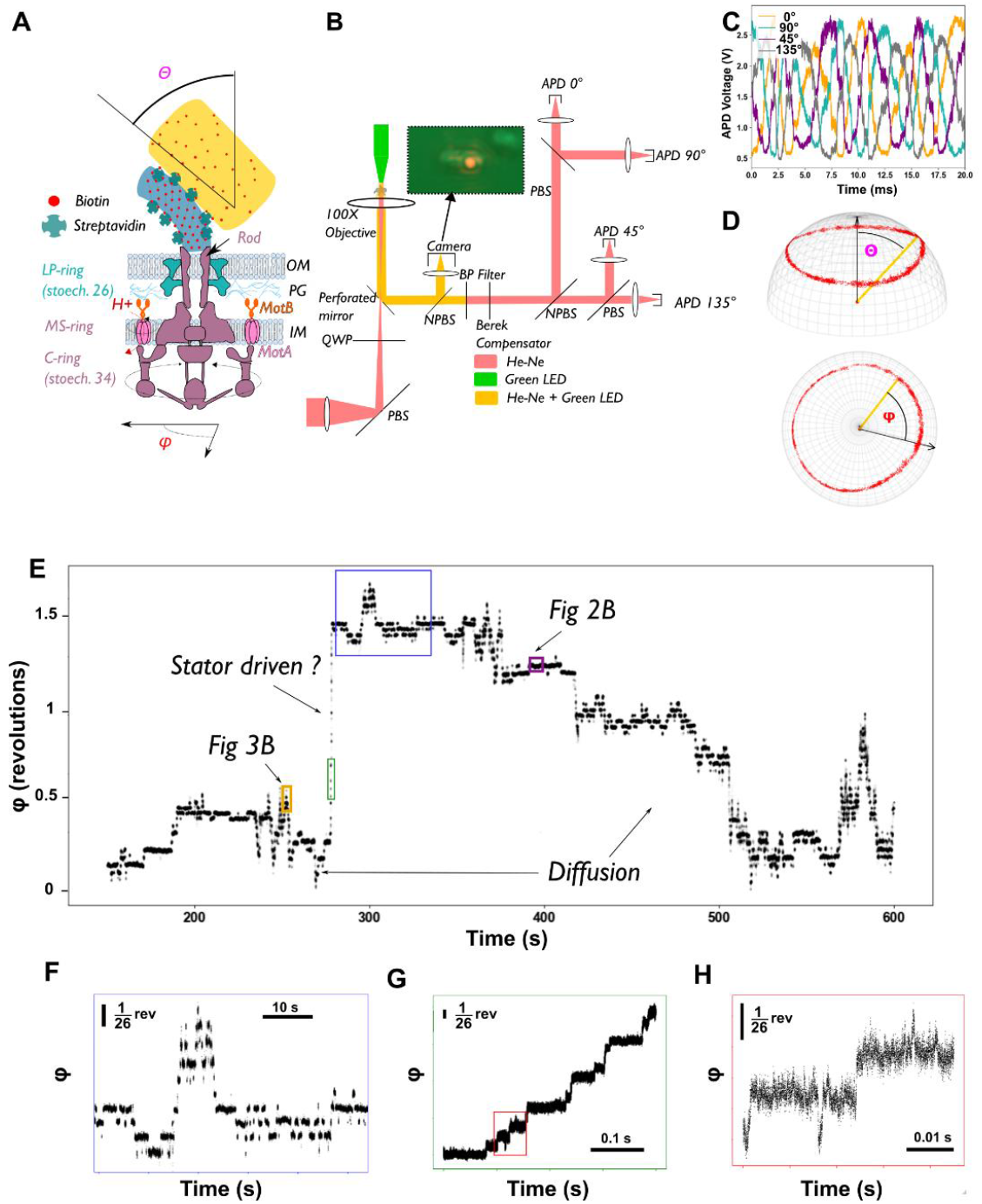
**A**. Top: Schematic of the structure of the bacterial flagellar motor and attachment of the gold nanorod. The internal part of the BFM has been magnified 2-fold compared to the hook and the gold rod. **B**. Schematic of the optical setup. Laser illumination at 633 nm (HeNe) backscattered by a gold nanorod is split into 4 polarization channels, each imaged onto an avalanche photodiode (APD). Inset: True-colour image of a nanorod (red) attached to a hook on an *E. Coli* cell (green). **C**. Raw polarization signals obtained from a rotating motor. **D**. Projection onto the unit sphere of raw nanorod orientations from the polarization signals shown in C and definition of the azimuthal angle ϕ and of the elevation angle θ. Side and top views. **E**. Unwrapped angle ϕ versus time of a single motor containing Na^+^-driven PotAB stators, in ∼0 mM Na^+^. **F**. Magnification of the blue rectangle shown in E. The motor displays discrete jumps between stable states separated by ∼0.25 radians. **G**. Magnification of the green rectangle of E, during a single unidirectional revolution. **H**. Magnification of the red rectangle in G, displaying a main transition and short transient jumps. (In E and F, only one data point in every 100 is plotted.)

Figure 1D and Supplementary Movie 1 show the orientations of a nanorod calculated from the polarization signals, using analytical formulae of Fourkas^37^ that model the rod as a dipole emitter. The data are close to the ideal case of a dipole emitter rigidly attached to a flagellum rotating purely around a single axis, which would appear in Figure 1D as a circle in a plane perpendicular to the rotation axis. The observed deviations from this ideal may be due to flexibility in the attachment between the hook and nanorod, shifts in the rotation axis, imperfect polarization properties of the optics, possible scattering or circular dichroism induced by the cytoplasm of the cell and deviations from pure dipole emission by the nanorod. We show details of our system in Supplementary Figure S1 and Supplementary Text S1, different methods to partially correct optical effects in Supplementary Text S2, and characterize the measurement noise of fixed rods in Supplementary Figure S2. We provide information about the orientation reconstitution in Supplementary Text S3. Estimations of the effect of losing some back-scattered light in the hole of the mirror (Supplementary Text S4, Supplementary Figure S3), the heating due to the absorption by nanorods (Supplementary Text S5), and the torque exerted by circularly polarized light (Supplementary Text S6) suggest that all three are negligible. We also measured the rotational drag of rotating rods attached non-specifically to a glass surface and found them to be diverse (Supplementary Figure S4 and Supplementary Text S7).

### 2.2 In the absence of motor torque the bacterial flagellar motor diffuses across a finite number of predominant energy barriers

Figure 1E shows versus time the azimuthal angle ϕ of the nanorod of Figures 1C&D, attached to a BFM in a cell expressing chimeric sodium stator complexes, PotAB^38^. We selected only nanorods like this one, with a rotational axis parallel to the optical axis, ensuring that the polar angle θ remains nearly constant and that ϕ accurately tracks the motor’s angular position. After replacing the buffer (40 mM NaCl) with a sodium-free solution (∼0 mM NaCl) at t = 0, the motor rapidly slows down from ∼1000 Hz to a few Hz. A few minutes later, it transitions into diffusional motion (see Supplementary Figure S5, upper left panel), interrupted by a single unidirectional revolution at t = 290 s, likely caused by either the transient reactivation of a single stator complex energized by residual trace sodium or an alternative unknown mechanism. Closer inspection (Figures 1F,G) reveals that both diffusion and unidirectional rotation consist of jumps between discrete metastable states, typically separated by ∼1/26 of a revolution. Figure 1H shows the data in the red box in Figure 1G with an expanded time-scale, illustrating rich dynamics extending to scales below 1 ms and ∼1/26 rev (Supplementary Figure S6). We observed similar behaviours across all BFMs in the absence of active stator units. Our analysis was based on 960 million position samples collected over 64 minutes from six different motors, representing two distinct strains: one expressing the PotAB stator and measured in sodium-free conditions, and the other lacking stators entirely (ΔMotAB). Supplementary Movie 1 provides an overview of an experiment on another PotAB motor, capturing the transition from ∼100 Hz rotation (in 5 mM NaCl) to passive diffusive motion.

### 2.3 Anomalous diffusion on multiple timescales

Figure 2A presents the time-averaged MSD of the angle ϕ as a function of elapsed time τ for six passively rotating motors, each recorded for over 400 seconds. Additionally, data from a gold nanorod fixed to the glass surface is included as a control. At shorter timescales, the MSD increases slowly with τ, indicating anomalous diffusion, and cannot be fitted by an exponential relaxation. At longer timescales, the MSD increases more rapidly, approaching but not reaching a slope of 1, which would indicate normal diffusion. In contrast, a standard overdamped Langevin equation in a smooth periodic potential with well-defined curvature would exhibit an initial exponential relaxation followed by a linear increase in MSD with a slope of 1 at longer times. Figure 2B provides a schematic representation that helps interpret the MSD behaviour in Figure 2A. The long-term diffusion observed in Figure 2A corresponds to transitions across barriers, while motion within these barriers follows a sub-diffusive relaxation. The inset displays a typical time trace of the angle during a dwell (corresponding to the purple frame in Figure 1E).

**Figure 2:**
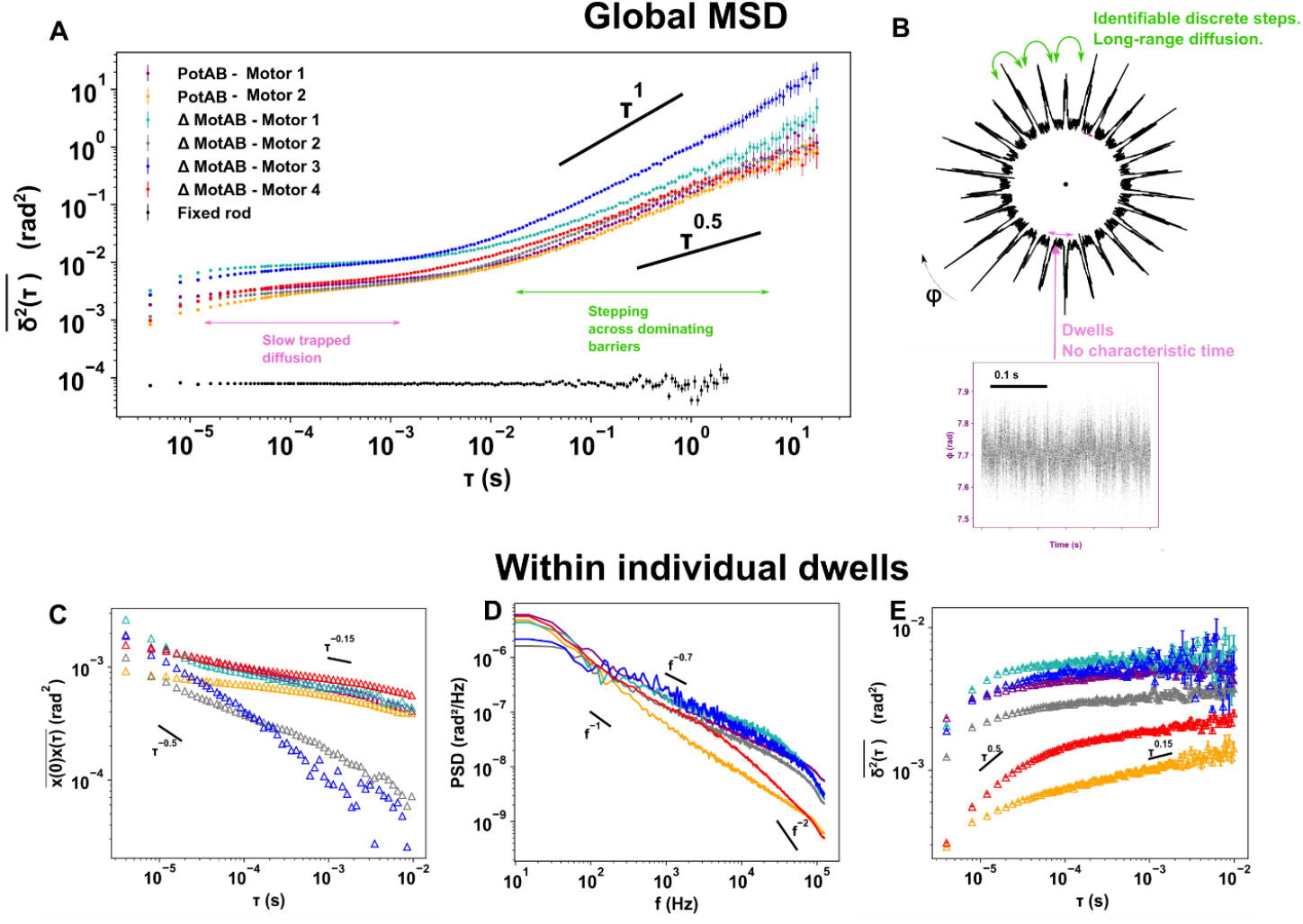
**A**. Time-averaged mean square displacement (MSD) of the angle ϕ as a function of elapsed time τ, for six passively rotating motors recorded for more than 400 s, along with a gold nanorod fixed to the glass surface as a reference. At short times, the motion exhibits slow anomalous trapped diffusion, while at longer times, a faster sub-diffusion is observed. In a simple overdamped Langevin equation within a periodic potential, one would expect a slope of 1 at long times and exponential relaxation at short times. Each point represents the average of all non-overlapping MSD values computed over full trajectories, with error bars indicating standard errors of the mean. **B**. Polar sketch illustrating the observed behaviour. The long-term diffusion observed in Figure 2A corresponds to transitions across barriers, while motion within these barriers follows a sub-diffusive relaxation. **Inset**: An angular time trace of a dwell (corresponding to the purple frame in Fig. 1E). **C**. Time-averaged autocorrelation function for six individual dwells from six different motors, with values averaged over 0.1 s. **D**. Power spectral density (PSD) of the same dwells as in C. **E**. Time-averaged MSD of the same dwells as in C and D, with values averaged over 0.1 s. For C, D, and E, refer to the legend in **A** to identify the corresponding motor for each dwell. The motor of Figs 1, 3A-C is PotAB-Motor 1.

Figures 2C, 2D, and 2E provide a detailed statistical analysis of the angular movement during six individual dwells—pauses in the angular signal exceeding 0.1 seconds—selected from six different motors. While there is significant variability in the signals across different motors, all cases exhibit sub diffusive behaviour characterized by non-exponential temporal auto-correlations with exponents ranging between −0.15 and −0.5 (Figure 2C). The power spectral densities are non-Lorentzian, with exponents between −0.7 and −1 (Figure 2D), except above the APD cut-off (∼125 kHz), and the mean square displacements (MSD) display non-exponential growth with exponents between 0.15 and 0.5 (Figure 2E). These MSDs fit very badly to an exponential relaxation model but reasonably well to stretched exponential or logarithmic models, and the process remains Gaussian across all timescales (see for example Supplementary Figure S7). The lack of a characteristic relaxation time within the dwells suggests the involvement of multiple processes, which could result from conformational changes in the numerous motor proteins (more than 50) that shape the rotational energy landscape, generating a dynamic potential with long-range correlations.

### 2.4. Transition times between the main stable states stretch over five orders of magnitude

The high resolution and quantity of our data allowed us to resolve directly heterogeneities that are usually experimentally inaccessible in systems that display anomalous diffusion. All six motors that we analyzed displayed jumps between local states separated by ∼1/26 revolution (Supplementary figure S6), but the exact absolute positions of the local states were slowly varying in time. The example in Figure 1 was an exception, with fixed state positions allowing us to define states and transitions consistently over the whole record. Defining steps is statistically challenging when the noise in between two steps spreads over so many time scales. To ensure consistency, we defined states that corresponded to local maxima in the histogram of angular positions over the whole record. Figure 3A shows the histogram for the motor of Figure 1, the only motor for which all angles were well sampled, displaying 26 stable peaks. To extract discrete transitions, we applied kernel-based change-point detection^39^ and only kept the change-points corresponding to transitions between states identified in global histograms (see Material and Methods). Figure 3B shows the section of data marked by the brown rectangle in Figure 1E, filtered in this way, illustrating that transitions separated by a few 10s of μs are well resolved. Each level of the red line corresponds to a peak identified in figure 3A.

**Figure 3:**
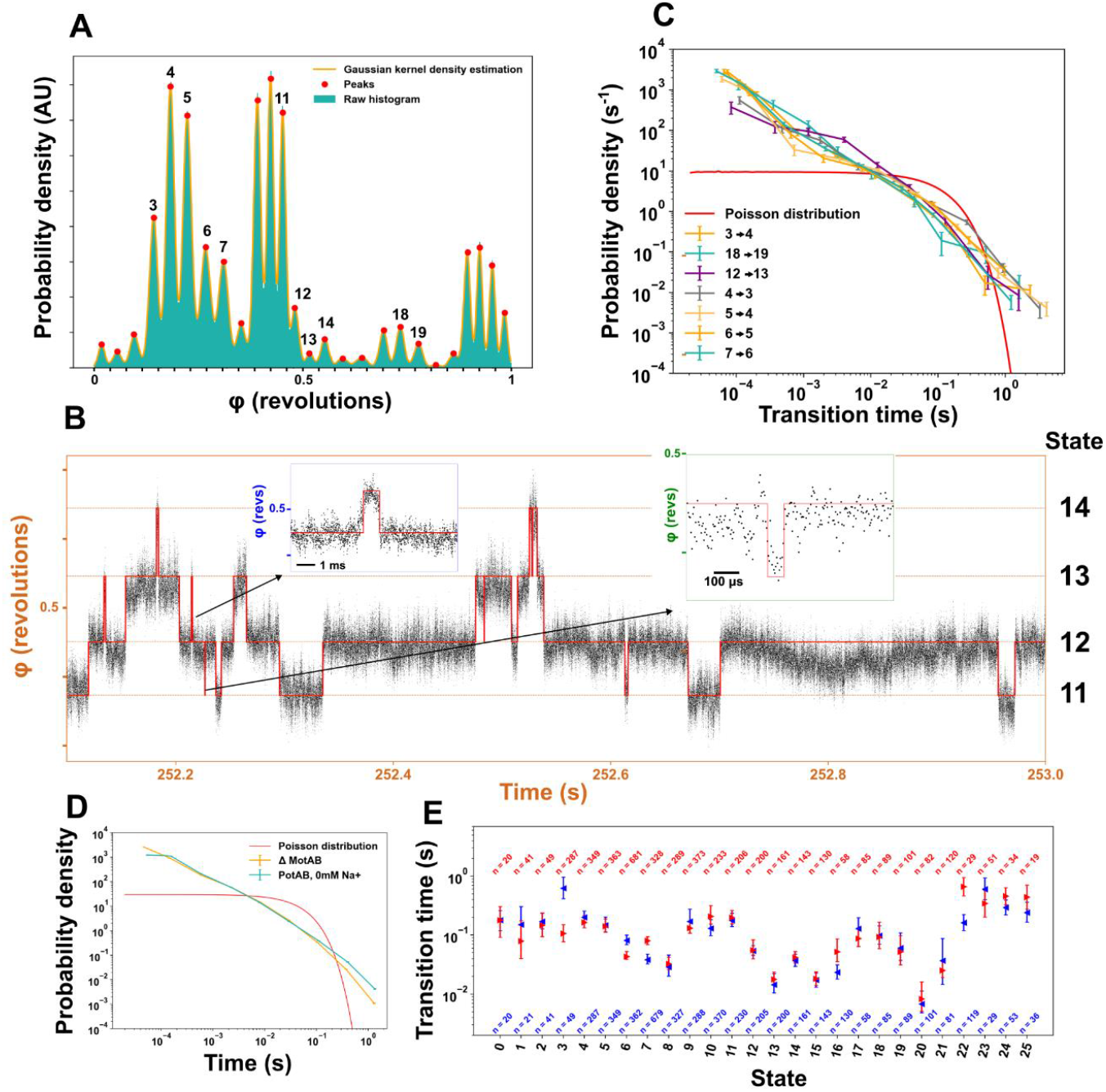
Transition dynamics between discrete states. **A**. Angular distribution (histogram and Gaussian kernel density estimation) of the whole trace shown in Figure 1, 26 local peaks (red) are used to define states. Tick spacing: 1/26^th^ of a revolution. **B**. Magnified portion of the trace in Figure 1 superimposed with the result of step fitting using the states identified in A as prior for the state position. Dwell times of order ∼0.1s (main figure), ∼1 ms (blue inset) and ∼50 μs (green inset) are all clearly resolved. **C**. Time distributions of seven transitions from states labelled in A to a neighboring state. Relative errors are computed as 1/sqrt(N), with N the number of transitions observed in each bin. Red: exponential distribution with the same mean as the distribution of times from state 3 to 4. **D**. Time distributions of all transitions between discrete states for all motors, either with no stator proteins (ΔMotAB) or with chimeric Na^+^ stators (PotAB) but sodium removed (the same motor as Figure 1 and 3A-C). Here all transitions are pooled since for most motors, one cannot define stable states over the whole traces. **E**. Average transition times out of each state, i, defined by a peak in A (blue: state i to state i-1, red: i to i+1). Error bars are obtained by bootstrapping 100 times each transition time distribution.

We define individual transition times 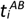 from one state (A) to another (B) as the total time spent in state A between transitions A -> B. For a Poisson process (no memory) the average transition time is the reciprocal of the transition rate *k*^*AB*^, the probability per unit time of jumping from A to B, and transition times are exponentially distributed. By contrast, each of the 7 best-sampled transitions showed an approximately power-law time distribution ranging from ∼10 μs to several seconds, with exponents varying between −1 and −1.5. (Figure 3C). Figure 3D shows the distribution of all transitions observed in the six motors, grouped by strain. All transitions are pooled since on most motors, most stable states are not fixed and can only be defined locally. The red lines in Figures 3C, D show very poor best fits to exponential distributions that characterize the classical barrier crossing Poisson process of Eyring and Kramers theories. Figure 3E shows the averages and corresponding bootstrapped errors for all transition times between all states identified in Figure 3A. Averages range from 0.006 s to 0.7 s, with some correlation between nearby states. This variation may represent static heterogeneity (stable variability in heights of the energy barriers separating the 26 minima), dynamic heterogeneity (variations between states reflecting global variations in time of the barrier heights between states), or a combination of both. It is likely that similar phenomena contribute to the anomalous diffusion that we observed in all motors studied (Figure 2A).

### 2.5 Transition rates and dwell positions vary with time

We directly observed two different types of dynamic heterogeneity in all motors measured: both the locations of the main angles and the rates of transitions between them changed on timescales of several seconds. Figure 4A shows motor angle versus time for one of the motor and Figure 4B the positions of peaks in dwell angle histograms of running data windows 10 s long, starting at 4 s intervals, versus the time at the beginning of the window. The inset to Figure 4B, reproduced in Supplementary Figure S8, shows histograms for two 10 s intervals separated by 40 s, highlighting a typical example of a transient change in the energy landscape. Before the change (red) the motor jumps between 7 angles, most of them ∼1/26 rev apart, afterwards (black) there are more peaks in the histogram, more closely spaced and at different positions. Similar shifts are visible in all motors observed (Supplementary Figures S9, Supplementary Movie S2-5). Figure 4C shows the MSD at τ = 0.1 s, calculated over overlapping 3.2 s data windows, versus the time at the beginning of each window. Non-homogenous variation over >2 orders of magnitude, even when the same angles are sampled (compare for example t=120s, t=220s, t = 500s), indicates that there are also shifts in the rate of transitions between energy minima.

**Figure 4:**
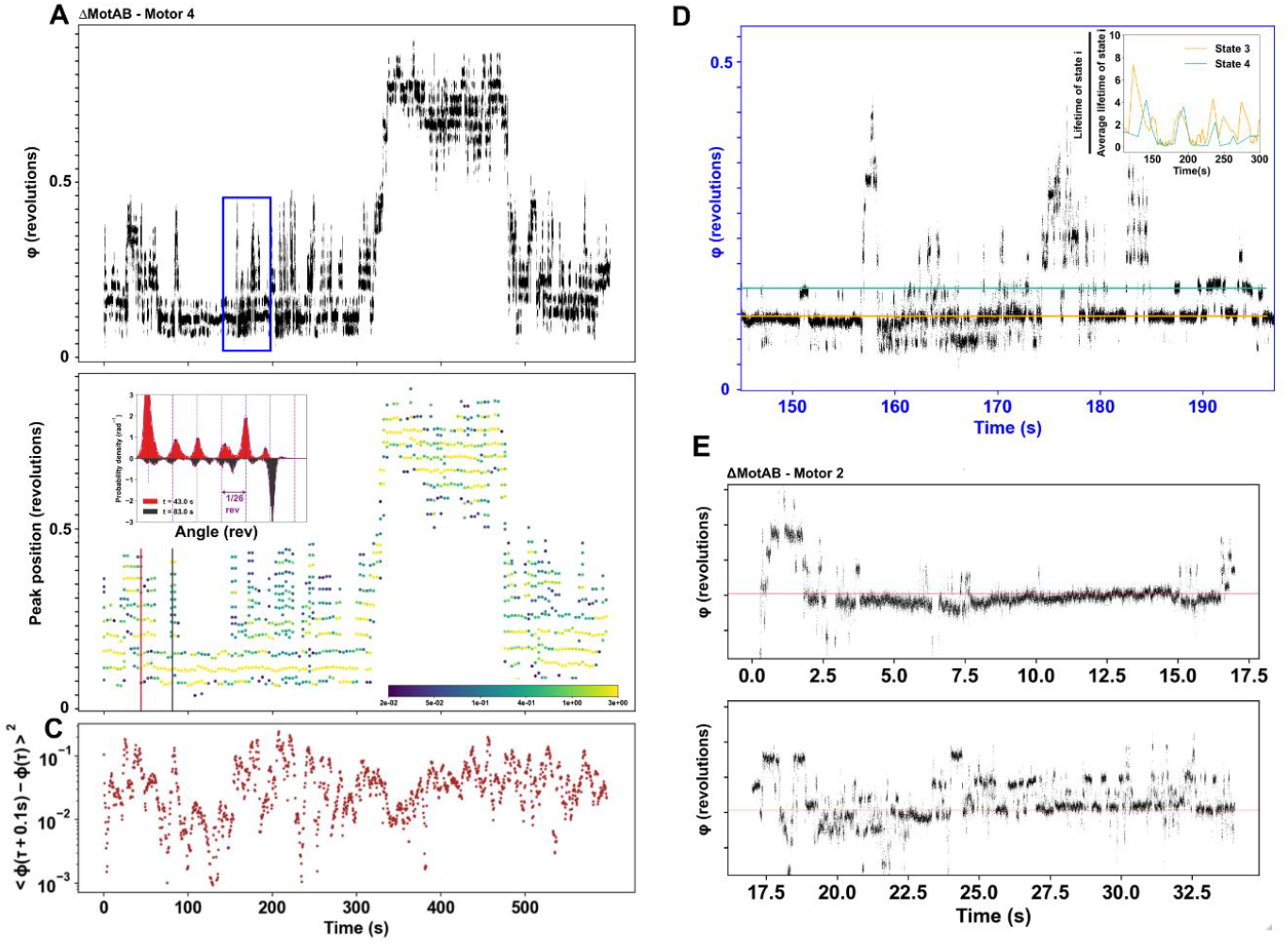
Heterogeneity of the diffusion dynamics. **A**. Angle *vs* time for a passively rotating motor. **B**. Evolution of the position of peaks in angle histograms of successive overlapping 10 s windows, starting at 4 s intervals. Color bar: probability density (rad^-1^) at each peak. Low peaks (dark blue) are more prone to errors. Inset: Angular distributions (histogram and Gaussian kernel density estimation) and peaks for 2 windows starting at times indicated by vertical lines in matching colours. Purple guide lines indicate 1/26^th^ of a revolution. See Supplementary Figure S8 for an enlarged version. **C**. Mean squared displacement at lag τ = 0.1s for overlapping 3.2 s windows in A, starting at 0.4 s intervals. **D**. Magnification of the blue rectangle in A. Horizontal lines mark angles of two states, identified by peaks as shown in B. Inset: Average duration *vs* average time, over 10 consecutive dwells in each state. Durations are divided by the average for each state over the entire record. **E**. Angle *vs* time for another passively rotating motor, showing abrupt change in the transition rates between the most occupied state (red line) and neighboring states. The lower trace immediately follows the upper.

Figure 4D zooms in on the data in the blue box in Figure 4A to examine in more detail the shifts in transition rate. The inset shows the moving average (over 10 neighboring dwells) of dwell times in each of two states marked by matching colours in the main figure, normalized by the averages over the full trace. The motor shifts between short-lived (∼ 0.005 s) and long-lived (∼ 0.5 s) transitions on a timescale of 10s of seconds. Some shifts alter the balance between the states (e.g. 250-300 s) while others do not (e.g. 180 s, 200 s). Figure 4E shows another example from another motor, also with stator proteins deleted (ΔMotAB - motor 2 in Supplementary Figure S9). Initially (upper) the state marked by the red line is favoured, with only short-lived excursions to the neighbouring states. Immediately afterwards (lower) the motor shifts: transitions are faster and neighbouring states now have similar occupancy. These two types of dynamic heterogeneity indicate that the energy landscape is shifting on multiple timescales, presumably due to some combination of shifts in the conformations of individual proteins, their relative arrangement in the bearing structure, or the rod sliding vertically in the bearing.

### 2.6 Symmetry and roughness of the energy landscape can be predicted from the atomic structure

Figures 5A-D show dwell angle histograms during powered rotation of one typical motor of each of 4 different types of BFM, either H^+^ or Na^+^-driven and clockwise (CW) or counterclockwise (CCW) rotating. All show 25-28 dominant peaks, consistent with previous measurements of flagellar rotation^29^ and with the atomic structure^5^ and passive rotation of the bearing. Random variability between motors, which was also observed previously^5,29^ is consistent with the observation of a highly dynamic and heterogenous bearing in our passive rotation experiments. Supplementary Figure S6 shows the histogram of all angular separations between adjacent dwell histogram peaks detected in all 6 passively rotating motors. The peak at ∼1/26 rev matches the previous results, and a second peak around 1/52 rev indicates either steps or phase-shifts of half the main step-size. The spread in peak separations may indicate variability in the LP-ring symmetry, but could also be due to limited success of our attempts at empirical correction of non-linearites in the measurement of motor angles (Supplementary Text S8).

**Figure 5:**
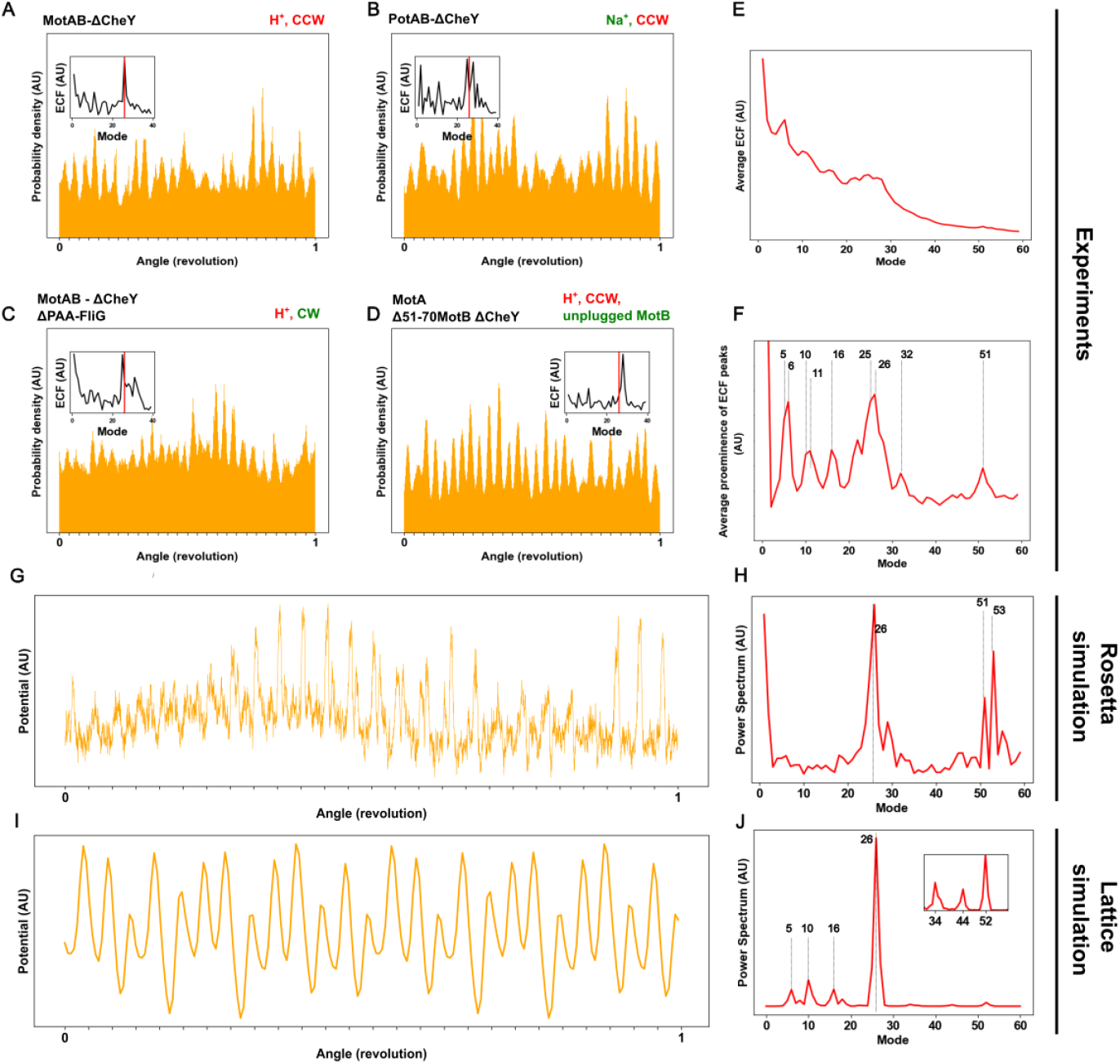
**A-D**. Histograms of the rotation angle ϕ averaged over 10s for four different motors of different strains. The variability of the histograms cannot be attributed to a particular mutant as within one strain they display similar variabilities. Insets: empirical characteristic function (ECF) showing the main periodicities in dwell angle. The red line shows the mode 26. CW: clockwise. CCW: counterclockwise. **E**. Average ECF over all non-overlapping 0.4 s windows (10^5^ data points) over 18 motors from different strains. The variation of background in the ECF from file to file smoothens the peaks. **F**. Prominence of periodicity in E. We define the prominence as the amplitude of a particular mode divided by the minimum amplitude over that mode and its four nearest neighbours. **G**. Energy landscape of the LP-ring / rod interface calculated via Rosetta using the cryo-EM structure from Salmonella (PDB 7CGO, (Tan et al. 2021), see Methods for details). **H**. Amplitude of the Fourier transform of G. **I**. Energy landscape simulated using a simplified lattice-model of the cryo-EM structure (Supplementary Text S10 and Supplementary Figures S10-14). **J**. Amplitude of the Fourier transform of I.

Figure 5A-D insets show the empirical characteristic functions (ECF, a spectrum of periodicities in the angle distribution^40^) for each motor, with peaks confirming the dominant periodicity near 26-fold. Figure 5E shows the average ECF over 18 different motors of all 4 types, and Figure 5F the prominence of ECF peaks at 5-6, 10-11, 16-17, 23-28, 32 and 51-52 per revolution, where again the spread of the measured periodicities may be due to non-linearitities in our measurement.

Figure 5G shows the energy landscape of the LP-ring / rod interface calculated via Rosetta using the cryoEM structure from Salmonella (PDB 7CGO^31^, see Methods for details), and Figure 5H the corresponding Fourier transform. The landscape is rough with 26 non-identical, highly anharmonic low energy “states” separated by variable barriers. Rosetta also predicts the observed ECF peak around 52. This may simply be a harmonic of the dominant 26-fold periodicity, but we note that the 1/52 rev separation of adjacent histogram peaks (Figure S6) cannot be explained this way. Rosetta does not predict the peaks that we observe at 5/6, 10/11 and 16, which may reflect the simulation’s very limited sampling of the conformational range of the bearing. These periodicities might also come from other parts of the motor. 11 notably is a sub-symmetry observed in the cryo-EM structure of the MS ring^41^. While Rosetta scores capture the relative magnitude and shape of the energy landscape given the atomic model^42^, its energy terms intrinsically approximate the distribution seen in the structures of the Protein Data Bank, and therefore cannot account for detailed transitions and kinetics of dynamic structures. Rosetta models also cannot, to date, efficiently computationally predict the experimentally observed variability, which could involve long range conformational flexibility.

Supplementary Text S10 describes a simple lattice simulation based on the arrangement of proteins in the rod. It predicts the observed bearing variability and most of the observed ECF peaks as arising from the symmetry mismatch between the 26-fold LP-ring and helical flagellar rod (Figures 5I, J). To reproduce the observed ECF peaks, the simulated interaction potential between each rod monomer and each repeating unit of the LP-ring must include features many times smaller than a single rod monomer, and all 26 LP-ring FlgHI units cannot be identical. Variability arises by simulating variation in the “defects” represented by non-identical LP-ring repeating units and/or by slight shifts in the alignment of rod and LP-ring.

## 3 Discussion

The atomic-resolution cryo-EM structure of the flagellar bearing^5,25^ is a compelling example of the possibility of atomically precise construction, a key goal in the emerging fields of nanotechnology and protein design. Our nanorod polarization method measures thermally driven passive rotation of single bearings with temporal and positional resolutions orders of magnitude better than has previously been possible. This reveals a reality very different from the static atomic perfection of the cryo-EM structure. Each bearing shows anomalous diffusion and structural dynamics over 4 to 5 orders of magnitude in timescale, characteristic of a system with a very large number of metastable configurations. While we cannot entirely exclude the possibility that external factors such as hook dynamics or cellular processes may contribute to the observed dynamics, the observed ∼26-fold symmetry is strong evidence that the bearing is the dominant contributor. Molecular processes characterized by multiple timescales and the absence of a single characteristic relaxation time has been observed in a wide range of cases; examples include the diffusion of particles in crowded and heterogeneous environments such as the cytoplasm^43–45^, protein folding in response to transient perturbations^46–48^, spectroscopic studies of populations of globular proteins^49^, spatial fluctuations in single complexes^50^, DNA hairpins^51^. We believe this to be the first direct observation of anomalous diffusion and its underlying structural dynamics in a passive, stably folded molecular machine. The function of the flagellar bearing, to transmit rotation from the motor in the cytoplasmic membrane to the extracellular filament, imposes no obvious requirement for structural dynamics of any kind. This raises the possibility that rich structural dynamics may be a universal feature of all large molecular machines.

Non-linear MSD vs time and non-exponential transition time distributions are typical experimental signatures of anomalous diffusion^52^. They indicate complex dynamics beyond the experimental resolution and are found in a variety of models featuring long-tailed energy landscapes^53^. Theoretical treatments postulate specific simplified models of this complexity and calculate predicted observable signatures for comparison with experimental data^52^. Our results at the shortest timescales (> ∼10 μs) within individual 26-fold dwells (Figures 2A-C), are an example of this typical case. MSD can be phenomenologically approximated by a stretched exponentials or logarithmic dependence on the lag, a behavior that has been called “ultraslow diffusion”^54^ and can be predicted by “comb” models^55^ of systems that can be trapped for highly variable durations in regions (the “teeth” of the comb) of a multi-dimensional space which allow no movement in the observable dimension. This fits with a picture of the flagellar bearing perpetually exploring a large multi-dimensional conformational space. Un-resolved dynamics within this space might correspond to fast rearrangements of small parts of component proteins, some of which would then be postulated to block rotation of the bearing. On intermediate timescales (∼0.01 – 1 s) we observed power-law transition-time distributions (Figure 3), that are most likely attributable to spatial and temporal heterogeneity in the interaction potential of the bearing^52,56,57^ caused by slower rearrangements of the protein structure. Our experimental resolution offers the prospect of direct observation of these heterogeneities, setting the challenge of developing new analytical tools to identify and characterize them. On the longest timescales (> 1 s) we observed unambiguous signatures of structural rearrangements (Figure 4), changing both the location of energy minima and the rates of transitions between them.

Our Rosetta simulations demonstrate that sensitivity to small features at the atomically-tight sliding interface of the flagellar bearing results in a rough interaction potential (Figure 5G). We speculate that the tight bearing interface might also be the key to our ability to resolve at the single-molecule level the rich dynamics that we observed, by generating large scale changes in the motion of the gold nanorod from small protein conformational changes at the interface. The current Rosetta based energy scoring algorithms cannot efficiently account for combinatorial explosion of the conformational space, which prohibits modelling of both static and dynamic heterogeneity. Instead, our simple model of the bearing symmetries shows that the sharp features predicted by Rosetta are necessary and sufficient to explain the observed periodicities in the bearing potential (Figure 5F), and that small shifts in these features can cause large-scale changes in the potential (Supplementary Figure S9).

What do our results tell us about molecular machines in general and molecular bearings specifically? First, that the flagellar bearing is smooth – it rotates under the influence of thermal fluctuations alone. The problem of making a smooth bearing from rough, discrete proteins has been recognized for nearly 50 years^6^. In 1978, Roger Hendrix^58^ proposed that symmetry mismatches, observed in bacteriophages, were an important ingredient in a smooth bearing - preventing high barriers or deep wells that would result if many proteins in the outer and inner rings could share the same high- or low-energy interaction state at the same time. The symmetry mismatches in the flagellar bearing fit this model. If we ignore the roughness and dynamics of the potential, which are likely to decrease the effective diffusivity^59,60^, we can use Kramer’s formula to estimate from the absolute transition rates an effective height of 11 – 14 k_B_T for the dominant 26-fold energy barriers (Supplementary Text S9), where k_B_ is Boltzmann’s constant and k_B_T the average thermal energy. A symmetric barrier *U* = 13 k_B_T high and *ϕ* = 1/26 rev wide corresponds to a torque of order *U*/*ϕ* = ∼200 pN nm, several times smaller than the torque generated by the flagellar motor^61^ and therefore smooth enough not to impede its rotation. This value is also comparable to the estimated torque per stator unit^62^, implying that the torque generated by a single stator and the energy barrier of the bearing may have coevolved to ensure that one stator can effectively overcome the bearing’s interaction.

Second, that protein machines can make perpetual thermal transitions within very large conformational state spaces, even when well folded, and even without any discernible functional requirement. This flexibility presumably allows for evolution to find sets of distinguishable sub-spaces that define the working cycles of molecular machines. For the flagellar bearing it is possible that conformational dynamics and a rough landscape may serve some function, perhaps contributing to easy rotation^15^. Alternatively, these may be universal default features of protein complexes, and there may simply be insufficient selection pressure on the bearing to remove them. Natural selection is always competing against genetic drift - unless selection pressure is strong, some non-optimized features may persist^63–65^. Future high resolution dynamical studies of natural and emerging designed^66^ molecular machines will reveal whether rich conformational dynamics are a universal and necessary feature of self-assembled macromolecular complexes.

## 4 Methods

### Attachment of gold nanorods to the hook of the bacterial flagellar motor

100 μL of overnight cultures of bacterial strains bearing AviTag mutations on FlgE^36^ are inoculated in 10 mL Tryptone Broth (10g NaCl, 5g Difco Tryptone) + 100 nM D-Biotin and grown at 30°C until they reach OD 0.4. 200 μL are then harvested and rinsed three times by successive centrifugations (2:30 minutes at 5000g) with 200 μL motility buffer (40 mM NaCl, 40 mM KCl, 10mM K_2_HPO_4_/KH_2_PO_4_ pH 7.1). 5 μL of neutravidin solution (90 μM in water) are added to the cells and we incubate 20 mins at room temperature while maintaining agitation. The cells are then rinsed 4 times with 200 μL motility buffer to remove excess streptavidin. They are centrifugated a fifth time and resuspended in 15 μL of motility buffer + 5 mg/mL BSA. 0.5 μL of biotinylated gold nanorods (Nanopartz C12-40-600-TB-DIH-50-1) are quickly added and the mixture agitated for one minute before transfer to centrifugal filtering columns with 0.22 μm pores (Millipore UFC30GV0S) and 150 μL of motility buffer + BSA are then added. To remove unattached gold, filtering columns are centrifugated 35 seconds at 5000 g (just enough to pass the solution but keeping the membrane wet). 200 μL of motility buffer + BSA are added again, and the filtering procedure repeated four times. For the last repeat, we do not centrifugate but instead vortex the column for one minute to detach the bacteria from the membrane. The solution above the filter is then retrieved, transferred to an Eppendorf tube, and centrifugated 2:30 minutes at 5000g, a smaller pellet should be observed with a slightly darker color, indicative of gold attachment. The pellet is resuspended in 50 μL motility buffer without BSA for later use.

### Microscopy setup

A Helium-Neon laser (12 mW, 633 nm, Melles Griot 25-LHP-991-230) is focused onto the back focal plane of an oil-immersion objective (100X Nikon Plan Fluorite Oil Immersion Objective, 1.3 NA, DIC) using a biconvex lens. A telescope made of two plano-convex lenses and two dichroic mirrors is added upstream to precisely adjust the focusing point of the laser. A linear polarizer and a λ/4 plate are added to circularize the excitation light. We ensure darkfield by drilling a hole (∼1 mm diameter) in a dichroic mirror at 45° to the optical axis, between the objective and the λ/4 plate, so that light reflected by the objective, coverglass and slide returns through the hole into the excitation path while light scattered by gold nanorods is mostly reflected into the emission path. The emission path consists of beam-splitters which separate the signal into four polarizations: 0°, 90°, 45°, and 135°. A first non-polarizing beam-splitter (Edmund Optics #35-963) reflects half of the light towards the camera. A second identical non-polarizing beam-splitter separates the light between two paths. On the first path a polarizing beam splitter with vertical axis separates the 0° and 90° polarizations. On the second path a polarizing beam splitter with an axis at 45° separates the 45° and 135° polarizations. A linear polarizer is added on each face of the polarizing beam splitters to remove leaks from s- and p-polarizations from the p- and s-faces respectively. Importantly, we use a phase compensator (in our case a Berek’s variable waveplate) to retrieve 45° and 135° polarizations: with the emission path in a horizonal plane, each dichroic mirror adds an uncharacterized phase shift between its s- and p-polarizations (0° and 90°). This does not affect the intensity of the 0° and 90° polarizations but mixes the 45° and 135° together. The Berek compensator (Newport 5540M) reverses the accumulated phase shift, un-mixing the 45° and 135° polarizations. The light in each polarization channel is then focused onto the capture area of one of four different avalanche photodiodes modules (Hamamatsu C5460-01).

### Cell strains

All strains used in this study are derived from the background strain *Escherichia coli* RP437. The hook protein FlgE contains an AviTag sequence (GLNDIFEAQKIEWHE) inserted between residues I221 and A222, “site C” as described by Brown *et al*.^36^. The AviTag is endogenously biotinylated by BirA.

Strains MTB32^67^ and MTB32-ΔCheY (MHR02) were derived from MTB9^36^. These strains lack the flagellar filament protein FliC and express MotAB natively. MHR08 is derived from MHR02 but contains a modified FliG in which the PAA169–171 segment has been deleted, as described by Togashi *et al*.^68^. This deletion results in clockwise flagellar rotation, consistent with prior studies in Salmonella enterica.

Strains MTB16, MTB24, and MHR05 are derived from YS34^29^, a strain with the following modification: ΔCheY, ΔMotAB, ΔFimA, and ΔFliC. MTB24 expresses the plasmid pYS13^69^, which encodes PotAB under IPTG induction. MHR05 carries the plasmid pMHR1, which encodes an unplugged variant of MotAB (MotB Δ51–70^70^). pMHR1 is derived from plasmid pDFB27^71^.

### Excitation path

We position the drilled mirror on a translation stage, the hole in the mirror being roughly aligned with the vertical optical axis. We then mount the objective on a threaded plate 2 cm above the mirror: it remains fixed throughout the experiment, focusing is performed by moving the sample vertically. The λ/4 plate and the linear polarizer (or a polarizing mirror) are placed below the drilled mirror. The focusing plano-convex lens is placed at its focal length from the back of the objective. A coverslip containing a dried solution of gold nanorods and micrometric beads is imaged on a camera using a lens of short focal length (5 cm) focused to infinity, and the sample moved vertically until beads appear sharp on the camera. Then the three following steps are repeated until convergence: 1. The laser beam is walked using two mirrors until it emerges vertically from the objective AND the area excited by the laser appears in the center of the field of view on the camera. 2. The lateral position of the drilled mirror is adjusted to maximize the intensity of the excited area and minimize scattered light at the mirror. 3. The position of the second lens of the telescope is adjusted to collimate the laser beam emerging from the objective. 1) and 3) are judged by observing the location and size of the laser spot on the ceiling.

### Emission path

All polarizing and non-polarizing cubes are positioned at the same height as the corresponding APD’s and carefully rotated – using the reflection of an alignment laser – so that their axis is perpendicular to direction of propagation of the light. A field of view containing a single excited nanorod is then used for fine tuning and the two following steps are repeated until convergence, to ensure that the gold rod is centered in the field of view and its image in the APD detector: 1. the orientation of the second non-polarizing cube is slightly tuned to maximize the signal in the first APD 2. the position of the nano-rod in the field of view is tuned using the nano-positioners to maximize the signal in the first APD. Then, the mirrors reflecting light to the three other APD are fine-tuned to maximize the signal in each detector. Note that to be able to correct their phase-shifts with the phase compensator, all dichroic mirrors should be placed in parts of the optical path where light scattered by the nanorod is collimated, since the induced phase shifts depend on the angles of incidence. The Berek compensator is aligned the following way. We first set its retardance to 0 and adjust its orientation so that the reflected beam propagates back along the incident beam. We then adjust the orientation of the plate so that when changing the retardance, the 0° and 90° APD signals remain unchanged. Finally, we used rotationally diffusing nanorods non-specifically attached to the glass to adjust the retardance of the plate to maximize the amplitude of variation of the 45° and 135° channels during rotation.

### Data acquisition

We recorded the voltage of each APD module (Hamamatsu C5460-01) at 250 kHz in differential mode using a multi-channel analog to digital converter (National Instruments PCI-6143) connected to the computer through a programmable chassis (National Instruments PXI-1071). The APD modules are connected to the converter through coaxial cables and a controller dock (National Instruments BNC-2090A). 100 kΩ resistors are soldered onto the R31, R32, R28, R27, R23, R24, R20, R19 slots of the controller dock, corresponding to the path between each analog input (+ and −) and the ground of each channel. This allows loading down the APD module output to the recommended range of 200kΩ as well as using floating sources: we power the APDs with two 12V batteries in series to avoid power-supply noise and earth loops in the circuit. APDs are insulated from the optical table. Black cardboard walls around them are coated with aluminum foil to shield the modules from electromagnetic noise (coming for example from the laser power units). The chassis is connected to the computer through a PCIe remote control interface card (51-924-001) on the computer side and PXI Remote Control Interface module (41-924-001) on the chassis side. The voltage of each APD module is recorded and streamed to file using a home-made LabView script.

### Flow cell preparation

A clean coverslip was first plasma-cleaned and assembled with a glass slide into a flow cell. The chamber was then incubated 5 minutes with a solution of 0.1M acetic acid and 0.015% chitosan (Sigma-Aldrich 448869) to promote bacterial attachment. The chitosan solution was thoroughly rinsed with motility buffer. After rinsing, cells were introduced in motility buffer and allowed to adhere. Non-attached cells were subsequently removed by gentle rinsing before imaging. Imaging was performed at 21°C in zero-sodium motility buffer (0mM NaCl, 80 mM KCl).

### Cell selection

The full 200 μm field of view is illuminated with a green LED (Thorlabs M530L3). We bring each bacterium one by one, with a manual nano-positioner, to the much smaller field of view excited by the laser (gaussian field of width σ = 1.4 μm). The maximum intensity of the laser excitation, calculated as 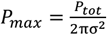, where P_tot_ is the full power measured with a power meter placed above the sample, is tuned to ∼200 W/cm^2^. A bright red point-spread-function is clearly distinguishable on the camera when a gold nanorod is attached to the bacteria. We display live the anisotropy of the polarization signal from the APDs as a function of time, that is 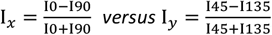 and the power spectral density (PSD) of I_*x*_ + iI_*y*_. The PSD of the combined anisotropy of a rotating rod attached to an active BFM displays a clear peak in frequency, indicating that the rotation is periodic. Displaying the anisotropies also allows us to differentiate between different orientations of the rotation axis and angle of rod attachment. Two similar overlapping circles in the space of anisotropies indicate a rod whose rotation axis is close to the optical axis - one circle corresponds to ϕ in [0,π], the other to ϕ in [π,2π]. When the rotation axis is further from the optical axis, two separate loops appear with different radii. We select the first type of trajectory to avoid points of low total intensity (rods near vertical, θ∼π/2), where the light mostly gets scattered perpendicularly to the optical axis and doesn’t enter the objective. At those points the optical noise is thus larger and makes the analysis harder to perform. By contrast, a rotation axis close to the optical axis allows us to consider the optical ϕ azimuthal angle of the rod to be equal to the angle describing the rotation of the motor, i.e., our variable of interest. These selection criteria reduce the throughput of the experiment in terms of cells. However, the throughput in terms of recording time is large due to the absence of photobleaching of the gold.

### Orientation reconstitution

The intensity of each of the four polarization paths focused onto APDs is measured at an acquisition frequency of 250 kHz. We reconstitute the orientation of individual nanorods using a previously published analytical formula for ideal dipolar emitters and perfect optics. Simple simulations based on ray optics suggest that the hole in the mirror should not affect much the inference of nanorod orientation, even though it removes part of the light scattered by the rod (Supplementary Text S4). There are two degeneracies affecting the orientation inference: the transformations ϕ->π+ϕ and θ-> −θ leave the polarization signal unchanged. The first is not problematic if θ is far from ±π/2 (rod orientation far from the optical axis) because we can unwrap ϕ by continuity. The θ degeneracy is not a problem either if the axis of rotation is close to the optical axis, so that most of the rotation of the rod is encoded in ϕ. We note that the degeneracies could be partially lifted by recording asymmetry in the scattered intensities in the back focal plane, which we have not attempted. To estimate the measurement noise due to the optics, we attach rods to the surface, infer ϕ and inspect the power density function of the signal. The ϕ signal of such a rod displays a standard deviation of ∼0.5° over a [1 Hz - 125 kHz] measurement bandwidth at a working APD voltage of 5V summed over the four APD’s.

### Molecular simulation of the interaction potential

We include the code for the Rosetta simulations at https://github.com/alexiscourbet/Bacterial-flagellum-energy-landscapes, along with the images referenced below. The structure of the bacterial flagellum was retrieved from the protein data bank (PDB: 7CG0, 7CGO_full1.png), aligned in xy to the cyclic symmetry axis, and truncated to only account for the molecular interactions at the interface between the stator and rotor (7CGO_trunc*.png). The rotational landscape was generated by rotating rod and LP-ring components relative to each other along the symmetry axis and sampling with a 0.1-degree angle (dock_rotor_axle_7CGO.py), producing a new structure for each conformation (/example_sampled_rotations/^*^.pdb). Rosetta energies were then computed for 3600 rotational bins x 10 trajectories for the whole rod and LP ring complex after repacking all residues (repack.xml). Inspection of the protein-protein interface of rotamer minimized structures between stator/rotor reveals low energy configurations and intricate hydrogen bonded interactions (7CGO_interfcaehbind1.png). The rotational energy landscape can be efficiently visualized as a polar plot showing the mean (+/-s.d.) (Figure 5G), with symmetric energy ‘spikes’ corresponding to high energies. A FFT representation of the computed energy landscape reveals dominant frequencies which correspond to experimental ECFs (Figure 5H).

### Data visualization and analysis

Large data files of tens of millions of points are smoothly visualized using a home-made program written in Python, available on Github (https://github.com/Mriv31/pyqtrod), which itself uses the visualization interface pyqtgraph (www.pyqtgraph.org). **Power spectral densities:** Power spectral densities (PSDs) are calculated using the Welch method from the `scipy.signal` module. A Hann window is applied with a 50% overlap. **Identification of global states:** The process begins with a smoothing operation applied to the unwrapped angles using a finite impulse response (FIR) window of size 100. Then a gaussian kernel density estimation (KDE) of bandwith 0.012 is applied to the wrapped angles. Peaks in the KDE curve are identified based on a prominence threshold of 0.01 (red points in Figure 3A), corresponding to global states that represent significant, recurring angular features. For most cells, we found that the “global” states are themselves not stable over the whole recording. They are thus calculated by applying the KDE estimation over 10 s windows. The positions of the identified peaks over the whole recording are shown as a function of time in kymographs such as in Figure 4B. **Identification of transition between states:** The method combines two complementary approaches to detect and refine state transitions in cyclic data: a local change point detection algorithm and the global state identification procedure described above. This combination allows for a sensitive detection of transitions while ensuring that the identified states are consistent and meaningful across the entire data trace. The process begins with identifying global states using the KDE method. Following this, a preliminary set of state transitions is determined using a step-finding algorithm, the Kernel Change Point Detection (KernelCPD) algorithm from the Python Ruptures package^39^. Given the absence of *a priori* knowledge of the statistical process underlying the data, the algorithm is configured with a linear kernel, a minimum segment size of 5 points, and a penalty value of 0.1, those parameters being adjusted empirically to ensure that there are more detected steps than are detectable by eye. The algorithm identifies the locations of change points, which are interpreted as the boundaries of discrete states. At this stage, given that the parameters have been chosen to give a high number of steps, some steps may be considered as spurious. These transitions are then refined by matching the detected states to the global states obtained from the KDE analysis. The matching process adjusts the mean value of the signal between two change points detected by the step-finding algorithm and attribute the corresponding part of the time trace to the closest global state. Overlapping segments are then merged. This approach strikes a balance: the local change point detection provides the necessary sensitivity to capture detailed transitions, while the global state identification ensures that these transitions align with the most significant and recurring features of the data. **Transition times and lifetimes:** Once the time trace has been reduced to a list of transitions between discrete states, we define the transition time from A to B as the time spent in A between two A->B events and we define the life time of A as the time spent in A between two events leaving A. **Empirical characteristic function:** The empirical characteristic function (ECF) characterizes periodicities in the angle distribution. The computation follows a bin-free sliding-window approach, dividing the signal into overlapping segments. For each window, the summation is computed over a window of 4 seconds (1000000 points). The magnitude of this sum is averaged across all windows to obtain the contribution of each mode to the ECF. Averaging the magnitude of the ECF computed over many different windows instead of performing a single ECF over the whole trace retains a periodicity even if its phase slowly drifts in time. In addition, we compute the “**local prominence”** of each peak among several cells. We define the local prominence of a mode in the ECF as 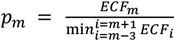. This prominence highlights the relative “sharpness” of the peaks compared to their local surroundings, which is particularly useful in mitigating the influence of noise-related variations in the background ECF signal.

## Acknowledgements

RMB and MR were supported by EPSRC Established Career Fellowship EP/S036660/1. MR was supported by the Human Frontier Science Program (fellowship LT0039/2022-C). ALN was supported by the ANR PHYBABIFO (ANR-22-CE30-0034) and PHYBION (ANR-21-CO15-0004) project grants. ALN is an Impulscience laureate of the Bettencourt Schueller Foundation, and this work was supported by Impulscience project PHYBION. ALN and AC were supported by the “Transatlantic Research Partnership”, a program of the FACE Foundation and the French Embassy. We thank David Baker and the Institute for Protein Design for providing resources and financial support for Alexis Courbet during the course of this research.

## Author Contributions

MR contributed to the design and implementation of the optical setup, instrumentation, and development of experimental protocols; conducted the experiments, developed the software, analysed and interpreted the data, created the figures and drafted the initial version of the manuscript. AN contributed to the design and development of the prototype for the polarization setup, participated in data discussions, provided the strain MTB32 ΔCheY, and contributed to the manuscript. AC performed the Rosetta simulations and contributed to the manuscript. HS contributed to strain design and modifications. RB designed the experimental setup, supervised the research, interpreted the first polarization signals, solved experimental roadblocks, discussed analysis and interpretation of the data, performed the lattice simulation, authored the majority of the final manuscript.

All authors discussed and approved the manuscript.

## Notes

### Competing Interest Statement

The authors have declared no competing interest.

