## Supplementary Information for "Single-molecule observation of multi-scale dynamic heterogeneity in the molecular bearing of the bacterial flagellum"

**Supplementary Texts S1 to S10**

**Supplementary Figures S1 to S14**

**References**

**Supplementary Movie 1:** Rotation of a PotAB motor as captured by our experiment. The motor is initially rotating at ~100 Hz in 5 mM NaCl and then goes into diffusion motion a few minutes after removing the sodium. Top left: Raw polarization data as a function of time. Top right: representation on the unit sphere of the inferred orientation ($\theta$,$\phi$) of the gold nanorod. Bottom left: Inferred angle $\phi$ as a function of time. Bottom right: Representation of $\phi$ on a unit circle.

**Supplementary Movie S2 to S7:** Dynamic representation of all the data presented in this work. Top: Positions of the preferred angular positions of the motor in each window. Middle: Windowed histograms of the angle $\phi$. Bottom: Enlarged view of the time trace of $\phi$. The red region corresponds to the window used to perform the histograms. **S2**: PotAB – Motor1. **S3**: PotAB – Motor 2. **S4**: MotAB – Motor 1. **S5**: MotAB – Motor 2. S6**:** MotAB – Motor 3. **S7**: MotAB – Motor 4.

### Supplementary Text S1: Alignment procedure

#### ***Main stage configuration***

The main stage is made of three horizontal plates placed directly in contact with each other.

The first, lowest level holds a micrometric translation stage supporting a dielectric mirror mounted at 45° to the optical axis, with a hole ~1mm in diameter along the optical axis, drilled by hand using a diamond drill bit.

The second level holds the fixed objective just above that mirror. The 45° mirror fits into the objective hole of that plate, in order to bring it as close as possible of the back focal plane (BFP) of the objective, which is inaccessible inside the objective structure for our typical 100X high NA objective.

That second level plate is also bored horizontally (~ 2 cm diameter). The space between the two first levels that is created by this cylindrical bore allows light reflected by the mirror to be transmitted horizontally into the excitation path.

The third, highest level holds the sample and its vertical and horizontal position can be adjusted with micrometric screws.

An opto-mechanical cage is mounted below the first level-plate. It contains a $\lambda$/4 plate and a 45° mirror reflecting the light from the horizontal excitation path into the objective.

#### ***Alignment of the excitation path***

*A. Preliminary alignment*

1. The drilled mirror is removed from the path. This allows the observation of the objective back reflection in the excitation path.
2. The objective is removed to allow a first approximate alignment of the laser. The latter is beam-walked until it goes through the centre of the hole holding the objective and reaches the ceiling of the room above the objective position.

3. The objective is placed back and will remain fixed. The laser is beam-walked with two mirrors until the back-reflection aligns with the beam.
4. The plano-convex lens ($f_0$ = 30 cm) is added to the excitation path at a distance $f_0$ +/- 0.5 cm from the objective exit pupil, using the back reflection to adjust its orientation and vertical position. It will then remain fixed.
5. A pair of lenses ($f_1$ = 10cm, $f_2$ = 20cm) are used as a telescope (separation $d = f_1+f_2$) to increase the laser beam width. A larger beam results in a larger aperture of the beam converging into the BPF of the objective and thus a larger excitation area in the field of view. Their vertical position and orientation are adjusted using their back-reflections.
6. The position of one of the three lenses, possibly held on a magnetic support or opto-mechanical stage, is adjusted until the beam exiting the objective appears at its smallest on the ceiling of the room (typically ~20 cm diameter). This focusses the laser beam into the back focal plane of the objective.
7. During steps 4-6, if all back-reflections remain aligned with the input laser beam, the laser beam should roughly exit the objective vertically. If it does slightly move laterally, adjust with the mirrors used to perform the beam walk in step 3.
8. Put the drilled mirror back in place. At first its body will appear red, because the laser beam scatters from the walls of the hole. Adjust its position until it becomes as dark as possible (some scattering may remain).
9. Adjust the axis of the $\lambda/4$ plate until the light exiting the objective is circularly polarized, or at least homogeneously polarized (the intensity of the light should not depend on the orientation of a linear polarizer placed between the objective and the power meter).

*B. Precise alignment of the drilled mirror.*

The drilled mirror will now be thoroughly centred with regard to the back focal plane of the objective.

1. Precisely assemble a camera at the back focal plane of a 4f lens setup to use as an imaging tool.
2. Image the back-focal plane of the objective with that setup (where the ceiling of the room appears sharp).
3. Put a paper in contact with the pupil of the objective, add a drop of immersion oil on the paper. Illuminate the full entrance pupil of the objective with a roughly focused LED. The paper serves as diffuser to fill the whole BFP of the objective. The back focal plane should now be a rather homogeneous circle with an unfocused dark central hole.
4. Slightly adjust the axial position of the imaging 4-f setup until the hole of the mirror appears sharp (typically a few mm). The back focal plane of the objective will still appear circular but slightly defocussed.
5. Adjust the lateral position of the mirror with the micrometric screws until the hole is centred compared to the circular back focal plane.

*C. Precise alignment of the excitation area*

The area of the field of view which is excited by the laser should now be centred so that all polarizations and rays scattered by the excited gold nanorods are captured homogeneously.

1. Assemble a camera at the focal plane of a short focal-length lens (~ 5 cm) to be able to observe the whole field of view of the objective.
2. Place the camera as close as possible to the lateral output of the stage, to image the full uncropped entrance pupil of the objective.

3. Prepare a coverslip with a dried solution of gold nanorods. Add a second coverslip, a thin paper and a drop of water on top. Place the sample on the sample stage.
4. Simultaneously illuminate the field of view from above, for example with a green light, as in B.3, and with the laser.
5. The gold nanorods will now appear in red in the excited area while the rest of the field of view will appear green. Due to the presence of the drilled mirror and the imperfection of the diffusion of the green light, the centre of the field of view might appear darker.
6. Adapt the height of the sample stage so that the gold nanorods are focused.
7. Beam-walk the laser beam until the excitation area lies in the centre of the field of view and the reflection on the sides of the field of view are minimal, as in Figure 1.

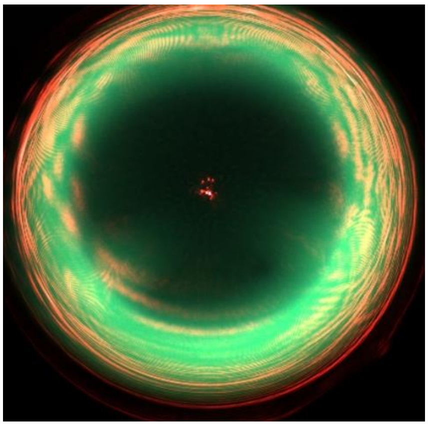

*An image of the full field of view after having centred the excitation area.*

***<u>Alignment of the emission path</u>***

*First, note that no lens should be placed between the objective and the horizontal mirrors preceding the polarizing cubes. Indeed, dichroic mirrors add a phase between s- and p- polarization which will affect the measurement of 45° and 135° polarizations and is strongly <u>DEPENDENT</u> on the incident angle. Any beam reflected before the separation of the polarization should be collimated so that all incident angles on the reflection area are equal.*

*<u>A. Preliminary alignment</u>*

1. Precisely measure the reflection and transmission coefficients of all non-polarizing and polarizing cubes of the setup, for both p- and s- polarized light. They will be used later to compute their effect on the measured signal.
2. Place a first non-polarizing cube path to send half of the light (reflected) to a bright-field camera placed in the back focal plane of a lens of focal length ~15-20 cm.
3. Take a sample with a low density of gold nanorods, and move the sample in the field of view until a single gold nanorod appears in the area excited by the laser. Focus the sample so that the nanorod appears sharp in the camera. Now, in a dark room and with the power of the laser at its highest, you can see a collimated image of the gold nanorod transmitted by the cube. It is circular (typically 5 mm of diameter with a 100X objective) with a hole in its centre, due to the drilled mirror. Use this collimated image to align the components of the emission path.
4. Place the first mirror about 1 meter away from the objective exit; this removes most of the spurious reflections by the objective, which are not collimated.

5. Place the second non-polarizing cube after the mirror.
6. On the reflecting side of the non-polarizing cube, place a polarizing cube to separate 0° and 90° polarizations. On the transmitting side of the non-polarizing cube, place a polarizing cube rotated by 45° to separate the 45° from the 135° polarization.
7. Add four lenses of focal lengths 50 cm and focus by eye the four images of the rod into the four different APD's.
8. Add the Berek compensator and the band-pass filter before the first mirror, making sure that the whole collimated image of the nanorod is centred with regards to their aperture.
9. After having roughly aligned the mirrors and cubes using the collimated image of a rod, add two pinholes aligned with the beam axis. Add a small alignment laser beam going through those pinholes and use the back-reflection of the cubes and lenses to precisely adjust their orientation.

*B. Fine adjustment of the mirrors and APD's position.*

1. Acquire the voltage of each APD.
2. Roughly position the illuminated gold nanorod in the centre of the excited area.
3. Adjust the position of the first APD until the signal is maximized.
4. Adjust the position of the gold to maximize the signal.
5. Iterate 3. and 4. until convergence. This ensures that the first APD is aligned with the position of maximum excitation in the field of view.
6. Adjust all mirrors reflecting the three other channels into the APD until each APD displays its maximum signal.

*C. Adjustment of the Berek compensator.*

The Berek compensator corrects the phase shift between s- and p- polarization which is introduced by the dichroic mirrors and affects the measurement of the 45° and 135° polarization.

1. Adjust the orientation wheel of the Berek compensator until rotating the dephasing wheel does not affect either the 0° or the 90° polarization signal. Fix the orientation wheel.
2. Find a freely rotating rod on the glass surface, they are typically non-specifically attached by the tip and display fast Brownian rotation.
3. Rotate the dephasing wheel of the Berek until the amplitude of variation of the 45° and 135° signals scattered by that rod become maximum. Note the position of the dephasing wheel.
4. Repeat step 3 ~10 times for 10 different freely rotating rods, the positions of the dephasing wheel obtained at each step should be very close. Average all measured values and fix the dephasing wheel at that position.

**Supplementary Text S2: Optical corrections due to cubes imperfections**

Each polarizing and non-polarizing cube has non ideal properties which affect the reconstitution of the polarization signal. To avoid any leak of the wrong polarization from each face of the polarizing cube, we glued linear polarizers on the exit face of each cube, aligned with the exit polarization corresponding to that face.

Then, the transmission of each polarization is carefully measured through the whole optical path by repetitive measurements and then averaging. We call $t_0$ the intensity transmission coefficient of a linearly polarized light through the whole 0° path and $t_{90}$ for the whole 90° path.

For the 45° and 135° path, the effect of the cubes is made slightly more complicated by the coupling between all polarizations. With $a$ and $b$ the *amplitude* transmission coefficient of the 0° and 90° polarization through all optical elements until the 45/135 polarizing beam cube, which are taken as real because their relative phase is cancelled by the Berek compensator, with $t_{45}$ and $t_{135}$ the *intensity* transmission coefficients through the 45/135 polarizing cubes of each polarization, and finally with $\alpha = \frac{a+b}{2}$ and $\beta = \frac{a-b}{2}$, we find that the relation between the incoming polarization intensities $I^{true}$ and the measured polarization exiting the series of cube $\mathrm{I}^{\mathrm{mes}}$ is:

$$\overbrace{\begin{bmatrix} I_0^{meas} \\ I_{90}^{meas} \\ I_{45}^{meas} \\ I_{135}^{meas} \end{bmatrix}}^{I^{mes}} = \overbrace{\begin{bmatrix} a_0 & & & \\ & a_{90} & & \\ & & a_{45} & \\ & & & a_{135} \end{bmatrix}}^{D} \overbrace{\begin{bmatrix} t_0 & & & \\ & t_{90} & & \\ -\alpha\beta t_{45} & \alpha\beta t_{45} & \alpha^2 t_{45} & \beta^2 t_{45} \\ -\alpha\beta t_{135} & \alpha\beta t_{135} & \beta^2 t_{135} & \alpha^2 t_{135} \end{bmatrix}}^{A} \overbrace{\begin{bmatrix} I_0 \\ I_{90} \\ I_{45} \\ I_{135} \end{bmatrix}}^{I^{true}}$$

The matrix A represents the imperfection of the cubes while the matrix D accounts for the effect of the APD's.

Then we compute $I^{true}$ from $\mathrm{I}^{\mathrm{mes}}$ by inverting this relation:

$$I^{true} = A^{-1}D^{-1}I^{mes}$$

While $A$ is measured experimentally and corresponds to the cube properties, $D$ is optimized by fixing $a_0$ to 1 and by minimizing the quantity $(I_0^{true} + I_{90}^{true} - I_{45}^{true} - I_{135}^{true})^2$ over a measurement of a nanorod in focus. This optimisation ensures that $I_0^{true} + I_{90}^{true}$ is as close as possible from $I_{45}^{true} + I_{135}^{true}$, which is a physical constraint. Note that this post-processing not only corrects the possible APD's biases but also any error of measurement of the $t_i$.

**Supplementary Text S3: Analytical inference of the rod orientation**

We use the formula derived by Fourkas, however we symmetrize their expression as a function of $I_0^{true}, I_{90}^{true}, I_{45}^{true}, I_{135}^{true}$ to use the information of all channels. In the following we drop the *true* index.

$$\phi = \frac{1}{2}\arctan 2\left(\frac{I_{45} - I_{135}}{2}, \frac{I_0 - I_{90}}{2}\right)$$

And for:

$$sin^2 \theta = \frac{4AI_s}{2I_d\big(sin(2\phi) + cos(2\phi)\big)\, C - 4I_sB}$$

, where *A, B, C* are constant defined in Fourkas' article, $I_s = (I_0 + I_{90} + I_{45} + I_{135})$, and $I_d = (I_0 - I_{90} + I_{45} - I_{135})$.

Due to degeneracies in the relation between the orientation and polarization signal, the angle $\phi$ is analytically retrieved between in the range [0, π] and θ in the range [0,π/2].

In order to get $\phi$ in the range [0,2π], we unwrap the signal so as to ensure continuity. This works without ambiguity as long as the nanorod angular trajectory does not come too close to the vertical, in which case we discard the data or only analyse the rotation speed by performing Fourier transforms of the raw intensities.

**Supplementary Text S4: Ray-optics simulation of the effect of a hole in the mirror.**

Since we cannot orient at will a gold nanorod, we do not have a simple experimental method to verify the accuracy of the orientation reconstitution from the four polarization channels. However, to get an intuition of how the hole in the mirror might impact the signal reconstitutions, we ran simple ray optics simulations corresponding to the analytical method of Fourkas. We use the far-field dipole radiation formula $\vec{E} \propto \vec{r} \times \vec{D} \times \vec{r}$, where **D** is the orientation of the rod and **r** the unit vector corresponding of a ray. We then attribute each ray to a position in the back focal plane (BFP). We add a mask in the BFP corresponding to the drilled hole in the mirror and integrate the four polarizations over the remaining BFP. Then we apply Fourkas' formula over those integrated polarizations and check that we retrieve the orientation of the originally defined **D(ϕ,θ).** Supplementary Figure S3 shows the result of the simulation of the dipole radiation as inferred in the back focal plane of the objective. It also shows the difference between the input orientations ϕ and θ, and the inferred orientation after integrating the intensities in the four polarizations in the BFP and applying on them the symmetrized Fourkas' formula indicated in Supplementary Text 3. The difference lies well below the degree (~0.001°), except in the case of a rod very close to the horizontal (θ~90°), which proves the correctness of the simulation and its agreement with Fourkas' analytical formula. We then add the Fresnel refraction at the water-glass interface and show that they don't have a very strong effect. We finally superimpose a mask corresponding to the true shape of the hole in our mirror onto the BFP before integrating. The simulation shows small errors compared to the noise of typical magnitude of the angular data observed in our data. Overall, while this is not an experimental proof, the simulations suggest that the hole in the mirror will have a negligible effect on the measurement and inference of the rod orientation.

**Supplementary Text S5: Estimation of heating due to the nanorod absorption**

The nanorods used in this study have diameter of 40nm and a length of 68nm. We can define their effective radius $\mathrm{R_{eff}} = \left(\frac{3\mathrm{V}}{4\pi}\right)^{\frac{1}{3}} \simeq 30\ nm$ and their aspect ratio $\frac{L}{D} = 1.7$. For those dimensions, Park et al. [1] have measured that the absorption cross-section of gold nanorods is of the same order of magnitude as the scattering cross-section, *i.e.* that at a given laser power, the nanorods will absorb as much heat from the laser as they scatter.

We can estimate the power scattered by the rod in our conditions. Roughly half of the light scattered by the rod goes into the objective, and half of that light goes to the APD's (the other to the camera). We work at a typical voltage, summed of the 4 APD's, of 4V, so that given the photoelectric sensitivity of the APD of $\epsilon = 1 \times 10^8 V/W$, we can estimate the total power scattered, and similarly absorbed, by one rod to be $P_{abs} = 0.15\mu W$.

To simplify, we model a gold nanorod as a sphere of radius a = 30 nm. In that case, the steady-state temperature profile around it is a solution of the 3D Poisson equation DT = 0, thus is equal to $\mathrm{T}(r) = T_{+\infty} + T_e \frac{a}{r}$. The total thermal flux, which corresponds in the steady state to the absorbed light is thus $4\pi T_e a\lambda = P_{abs}$, with $\lambda = 0.6\ W/m.K$ the thermal conductivity of water. Thus $T_e = \frac{P_{abs}}{4\pi\lambda a} \simeq 0.75\ K$.

Thus, the heating due to the nanorod absorption cross-section does not seem to be critical. However, one should be aware that either an increased absorption coefficient (which could be caused by a smaller aspect-ratio of an individual rod), or a slightly larger laser power would bring the temperature increase to a range of a few Kelvins. Caution should be exerted while choosing those parameters.

**Supplementary Text S6: Estimation of torque exerted by the circularly polarized light**

The optical torque exerted at the plasmon resonance for slightly larger rods has been previously estimated to lie [2] around $10\ \mathrm{pN.\mu m}/(W.\mu\, m^{-2})$. Our laser beam in focus has a Gaussian shape of deviation $\sigma$ = 1.4 µm (see figure below). The typical laser power used in our experiment (as measured with a power meter placed above the glass slide) is 40 µW. Thus, the maximum light intensity at the centre of the beam reaches 3 µW/µm$^2$. Using the estimation above, this corresponds to a torque of 0.03 pN.nm, which is negligible in comparison with the typical torque per stator of the bacterial flagellar motor of 200 pN.nm [3]. Nonetheless, based on the rotational drag measured from the rotation of nanorods non-specifically attached to the glass (Supplementary Figure S4 and Supplementary Text S7), we estimate that circularly polarized light should bias the rotation of unhindered rods by approximately 10 revolutions per second. This is somewhat greater than the bias observed in non-specifically attached rods exploring the full angular space, which is a few revolutions per second.

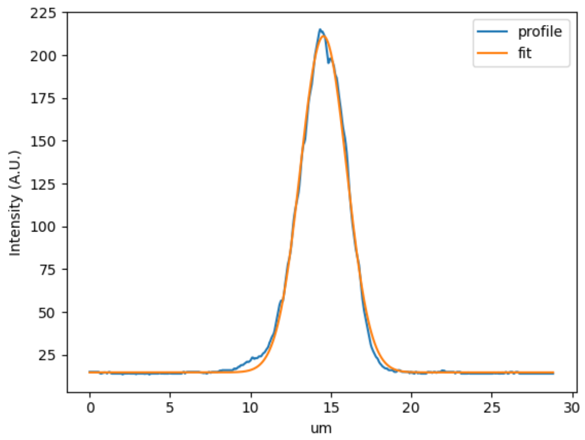


**Supplementary Text S7: Rotational drag of rods non-specifically linked to a glass surface.**

We measured the rotational drag of ten gold nanorods which were undergoing Brownian motion while being non-specifically attached to the glass surface. We expected that the PSD of ϕ would decrease as $\frac{1}{f^2}$, with f the frequency. We show their power spectrum density in Supplementary Figure S4. Over the ten gold nanorods, six display such an expected spectrum while four others seem partially trapped at a preferred angular position and only display the -2 exponent at high frequency, a region which we use to infer the drag. We find that drag values accumulate around $6.10^{-4}\ pN.nm.s.rad^{-1}$, however with a large variability, some rods displaying drags as large as $1.10^{-2}\ pN.nm.s.rad^{-1}$ . Series development of the rotational drag based on boundary element computations[4,5]predict the rotational drag of our 68x40 rods to be equal to $\gamma_{\perp} = 5.4\ 10^{-4}\ pN.nm.s.rad^{-1}$, which is the same order of magnitudes as our smallest experimental values. As a comparison, previous studies used 60nm[6] and 100nm[7] gold nanospheres, which total rotational drag coefficients are similar to the one measured here (typically from 0.001 to 0.005 $pN.nm.s.rad^{-1}$ ).

**Supplementary Text S8: Empirical attempts to correct non-linearities**

The orientation of a nanorod, $\phi_{measured}$, may contain systematic errors due to nonlinearities in the measurement process. To empirically correct these errors, we compute a correction function FF such that the corrected orientation, $\phi_{corrected} = F(\phi_{measured})$, restores the accuracy of the measurements over the periodic domain [0,2π].

**Correction Hypothesis and Approach**

The correction hypothesis is based on the assumption that the high-frequency (HF) noise in $\phi_{corrected}$ is isotropic, i.e., the HF displacements $\sigma = \phi_{corrected}[i+1] - \phi_{corrected}[i]$ should not depend systematically on $\phi_{corrected}$ . This isotropy is disrupted by nonlinearities in , $\phi_{measured}$, leading to systematic variations in measured displacements.

**To derive F, we proceed as follows:**

- **Bin and Measure Displacements:** The domain [0,2π] is divided into N equally spaced bins. The standard deviation of the measured displacements is calculated in each bin. This provides a binned measure of HF noise $\sigma(\phi_{measured})$.
- **Filter Noise**: To mitigate spurious effects of measurement noise and undersampling, a low-pass Fourier filter is applied to $\sigma(\phi_{measured})$.

- **Integrate for Correction**: The inverse of the filtered $\sigma(\phi_{measured})$ is integrated cumulatively to construct a monotonic mapping F.

Without prior assumptions about the periodicity of ϕ, the correction function F tends to amplify periodic patterns. This suggests that it effectively compensates for distortions introduced by the nonlinear measurement process. Visual inspection of the filtered and corrected data demonstrates improved alignment with the hypothesis of HF noise isotropy, providing qualitative validation of the method.

**Supplementary Text S9: Estimation of Barrier Heights Using Transition Times and Drag Coefficients**

Kramers' formula for transition rates over energy barriers in a smooth parabolic potential, mirrored at the peak of the barrier, is a widely used tool in biophysics to estimate energy barrier heights from transition times and drag coefficients. In our study, we estimated the rotational drag coefficient for gold nanorods by combining theoretical approximations and experimental measurements of nanorods attached unspecifically to a glass surface (Supplementary Text 7). The rotational drag was found to be approximately $\gamma = 6.10^{-4}\, pN.nm.s.rad^{-1}$albeit with significant variability due to differences in nanorod shapes.

Kramer's formula for a mirrored parabolic potential where two minima are separated by a distance δ is $t \propto \frac{\sqrt{\pi}\gamma\ \delta^2 \sqrt{T}}{4\, U^{3/2}}\ e^{U/T}$ [8], p. 293

For the typical transition time 0.1s found in this study between states separated by δ = 1/26 rev, we find an energy barrier of 14 $k_BT$. If we now take the largest drag coefficient measured on a freely rotating rod ($\gamma = 1.10^{-2}\, pN.nm.s.rad^{-1}$), we find an energy barrier of 11 $k_BT$.

Despite its utility, several limitations apply to the use of Kramers' formula in estimating barrier heights. Kramers' formula assumes a smooth parabolic potential near the barrier. This approximation greatly simplifies the relationship between the barrier height and the transition times. However, the exact shape of the energy landscape can significantly alter the dynamics. For instance, a squeezed barrier—where the potential narrows near the top—can increase transition rates by reducing the time spent near the barrier. Such deviations from the assumed parabolic shape lead to estimates that reflect an effective or equivalent barrier height rather than the true microscopic energy barrier. Furthermore, in a landscape accurately described by Kramers' model, transition times are expected to follow an exponential distribution. Our experimental data, however, exhibit a clear deviation from this distribution. This discrepancy indicates that the true energy landscape underlying the transitions is more complex than the idealized Kramers potential.

**Supplementary Text S10: Symmetry model of the Flagellar Bearing.**

We describe a simple simulation that makes the following predictions.

1. Most of the observed ECF peaks arise from the symmetry mismatch between the 26-fold LP-ring and helical flagellar rod.
2. To reproduce the observed ECF peaks, the simulated interaction potential between each rod monomer and each repeating unit of the LP-ring must include features many times smaller than a single rod monomer, and all 26 LP-ring repeating units cannot be identical.
3. The observed variability arises by simulating variation in the "defects" represented by non-identical rod or LP-ring repeating units and/or by slight shifts in the alignment of rod and LP-ring.

Fig. 5F in the main text identifies periodicities in flagellar rotation of 5, 6, 10, 11, 16, 32 and 51 in addition to the dominant periodicity around 26-fold. 5, 6 and 11 match the 5-, 6- and 11-start helices that characterize the protein lattices that span the distal components of the bacterial flagellum - from the export apparatus at the base of the rod, all the way to the tip of the filament. In particular, the flagellar bearing consists of the cylindrical rod which is a polymer built upon this helical lattice, rotating inside the 26-fold LP-ring which makes tight contact with the rod around one particular circular narrowing of the inside face of the LP-ring ([9] see Supplementary Figure S10 ).

We made a simple model of the interaction potential of the bearing to explore whether the symmetry mismatch between the rod and LP-ring might be responsible for the observed periodicities in rotation.

**Model**

The model places a rod monomer at each lattice site, approximately in the pattern of the Cryo-EM structure. Each rod monomer is represented (in cylindrical 2D co-ordinates representing the circumference and axis of the flagellar rod) by a small number of Gaussian potentials of interaction with each LP-ring repeating unit. Multiple Gaussians allow rod monomers to be asymmetric, and the inclusion of narrow gaussians permits "sharp", short-range, features to be modelled (Supplementary Figure S11).

Each LP-ring repeating unit contributes a 1-dimensional z-section of the rod monomer potential to the total bearing rotation potential (Supplementary Figure S12, top), at a z value corresponding to the axial position of the LP-ring (Supplementary Figure S11, purple line). This is motivated by the structure which shows a very narrow "collar" where the LP-ring is in atomic contact with the rod. The total bearing rotation potential (Supplementary Figure S12, middle) is the sum of all 26 individual LP-ring repeating unit contributions, each phase shifted by 1/26 of a rev. (The 26-fold LP-ring periodicity can be replaced by other values to model different LP-ring symmetries.)

**Results**

Supplementary Figure S12 shows the monomer potential, the total bearing potential and its Fourier power spectrum, for the rod potential and LP-ring axial position of Supplementary Figure S11, in the case where all 26 LP-ring repeating units are identical. The peak periodicities at 6, 10, 16, and 26 are all observed in the experimental data, as is a peak near 52 which we attribute in the model to a harmonic of the dominant 26-fold periodicity. The experimental peaks observed at 5, 11 and 32 are weak in this case.

Supplementary Figure S13 shows the effects of defects in the rod lattice and LP-ring. Removing the sharpest feature of the rod potential, present in Supplementary Figure S11 due to a unique "defective" rod monomer, greatly reduces the 26-fold peak (top). Adding a defect in the LP-ring, modelled here by reducing 5-fold the strength of the interaction potential with one of the 26 LP-ring repeating units, enhances the 5, 6, 10 and 11-fold periodicities (middle). These effects are additive (bottom). This is to be expected: a period-1 defect in the LP-ring probes the helical lattice of the rod, which would otherwise be smoothed out by the non-commensurate 26-fold LP-ring symmetry. Similarly, sharp features on the rod lattice are period-1 probes of the 26-fold LP-ring – they do not average because each rod lattice site has a unique position in relation to the tight constricting ring in the bearing.

Supplementary Figure S14 Illustrates that any sharp features at the bearing interface will generate 26-fold periodicity in the total potential – here they are present in all rod monomers of the majority type (FlgG-b) at the constricting ring. The details of the potential are extremely sensitive to the exact positions and shapes of these sharp features. However, a small peak at 34 (close to the experimentally observed 32-fold peak) appears under most parameter sets of the model.

We were unable to identify parameters that substantially enhance the peak at 16, but many parameters sets remove it or shift it to 18.

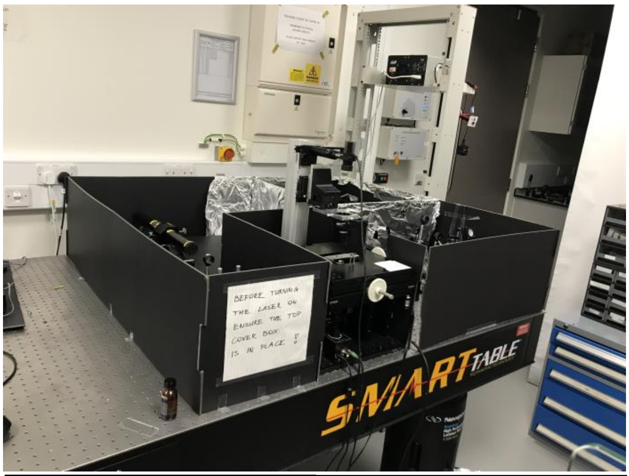


1. HeNe Laser 12 mW
2. 200 mm plano-convex lens
3. 100 mm plano-convex lens
4. λ/2 plate (633 nm)
5. 300 mm plano-convex lens
6. Polarizing beamsplitter
7. λ /4 plate (633 nm)
8. Drilled dielectric mirror
9. Non-polarizing beamsplitter
10. 150 mm plano-convex lens

11. Color camera
12. 633 nm band-pass filter
13. Berek's compensator
14. Polarizing beamsplitters
15. 500 mm plano-convex lens
16. APD modules

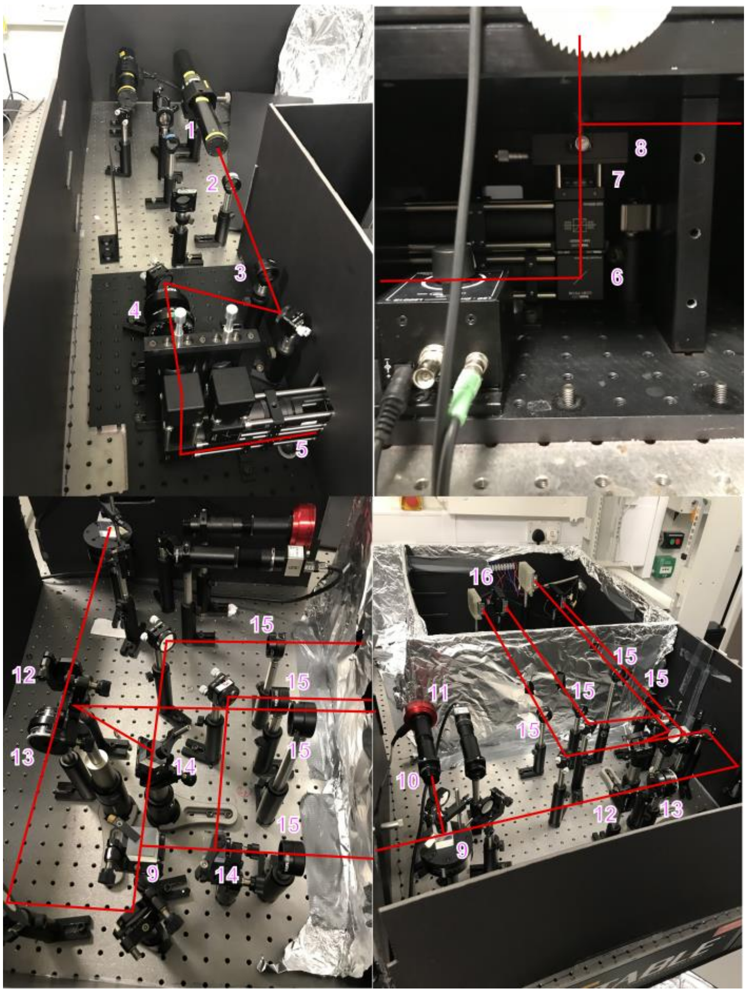


**Supplementary Figure S1:** Detailed pictures of the experimental setup: See Figure 1B and associated text for further descriptions of the components and their roles.

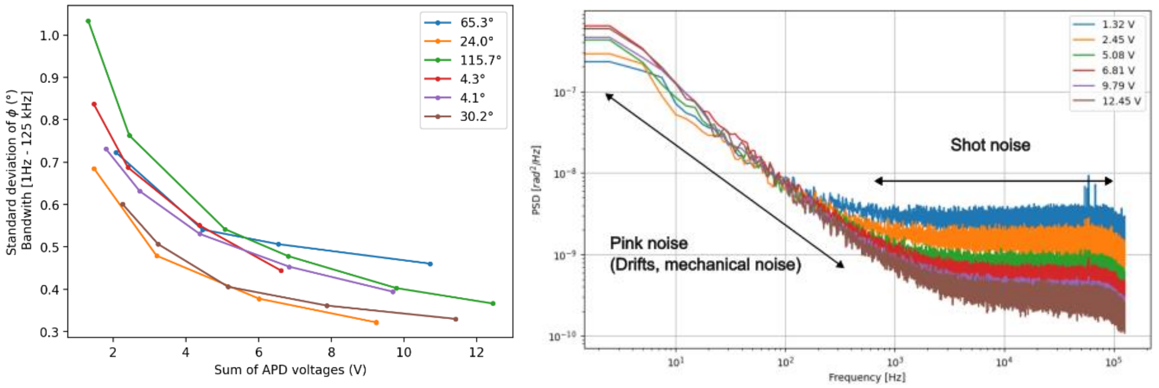


**Supplementary Figure S2:** Measurement noise of the orientation of five stuck rods on a glass surface. **Left**: Integrated noise in degrees as a function of the captured light intensity, modulated by changing the laser power. The labels indicate the average orientation $\phi$ of the rod. **Right**: Power spectral density of the angular signal of the stuck rod corresponding to the integrated signal in green on the left. The flat portion at high frequencies corresponds to the shot noise and depends on the light intensity received in the APD. The left portion, with exponent -1, is pink noise due to various drifts in the optical system.

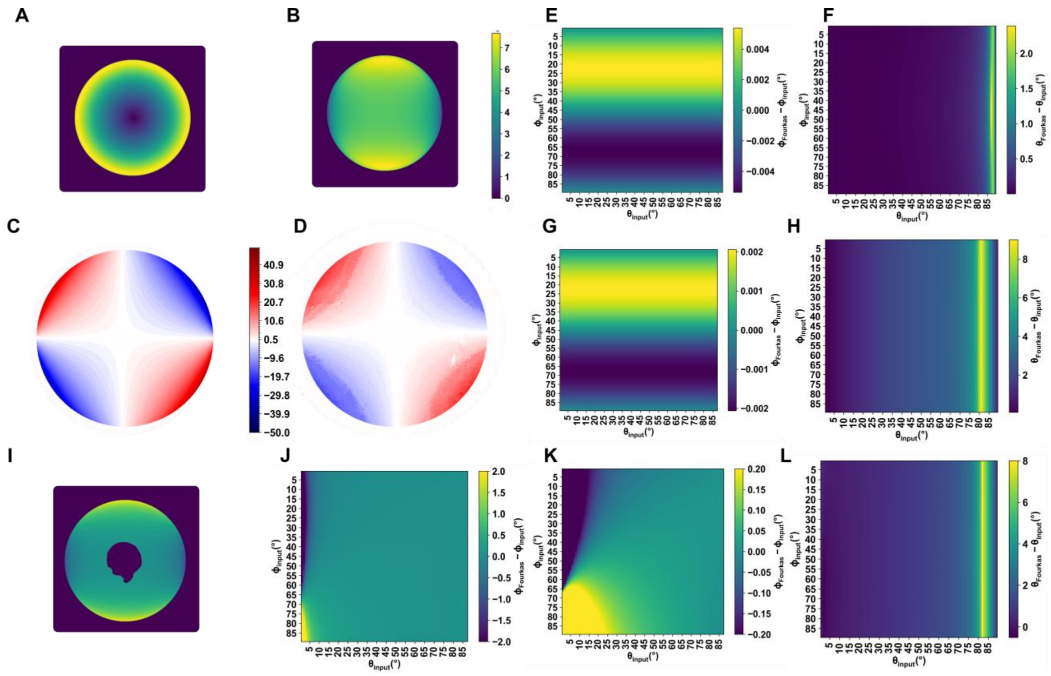


**Supplementary Figure S3: Simulation of the radiation of a dipole in focus as observed in the back-focal plane. A-B.** Intensity profile in the back focal plane (arbitrary units). **A**: $\theta$ = 0°, **B:** $\theta$ = 90°. **C**: Simulated orientation of the electric field as a function of the position in the back focal plane (in degrees). **D.** Polarization direction as measured in the back focal plane of the objective by putting a linear polarization filter in focus. **E.** Difference between $\phi$ used as input in the simulation and the retrieved angles after applying Fourkas formula to the integrated intensities in the back focal plane. As a function of $\phi$ and $\theta$ used as simulation input. **F.** Same as E. but for $\theta$. **G-H.** Same as E-F but adding in the simulation the effect of refraction on polarization at the water/glass interface. **I.** Same as B. but superimposing the hole in the mirror to the BFP. **J-K-L**. Same as E-F in the case of a hole in the back focal plane. **J** and **K** only differ by the angle range used in the colour bar. Deviations are less than 1 degree except in $\theta$ very near $\theta$= 90° (horizontal rods) and in $\phi$ very near $\theta$= 0° ( vertical rods ).

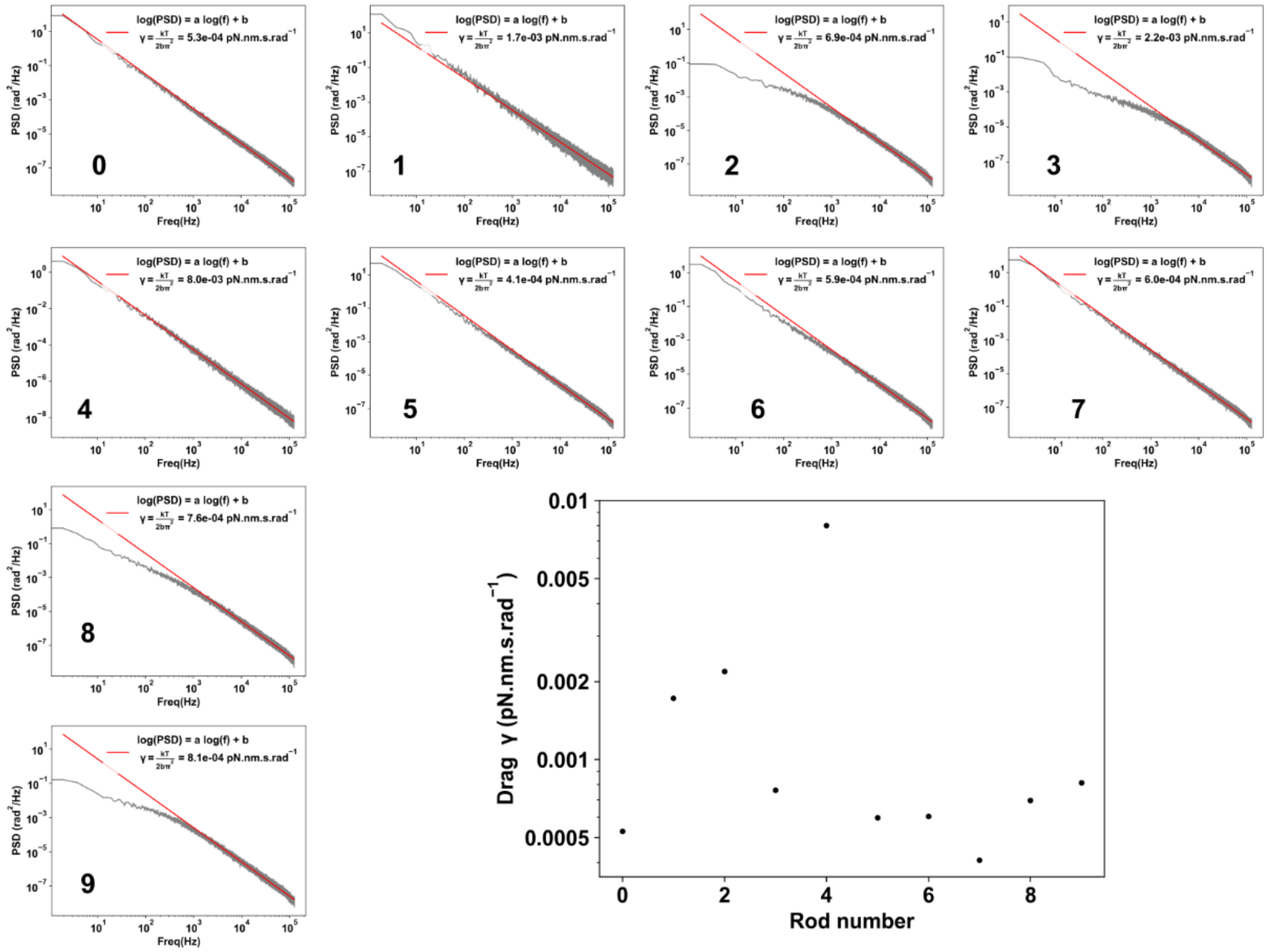


**Supplementary Figure S4: Power spectrum density of 10 different rotating rods on the surfaces and their inferred rotational drags.**

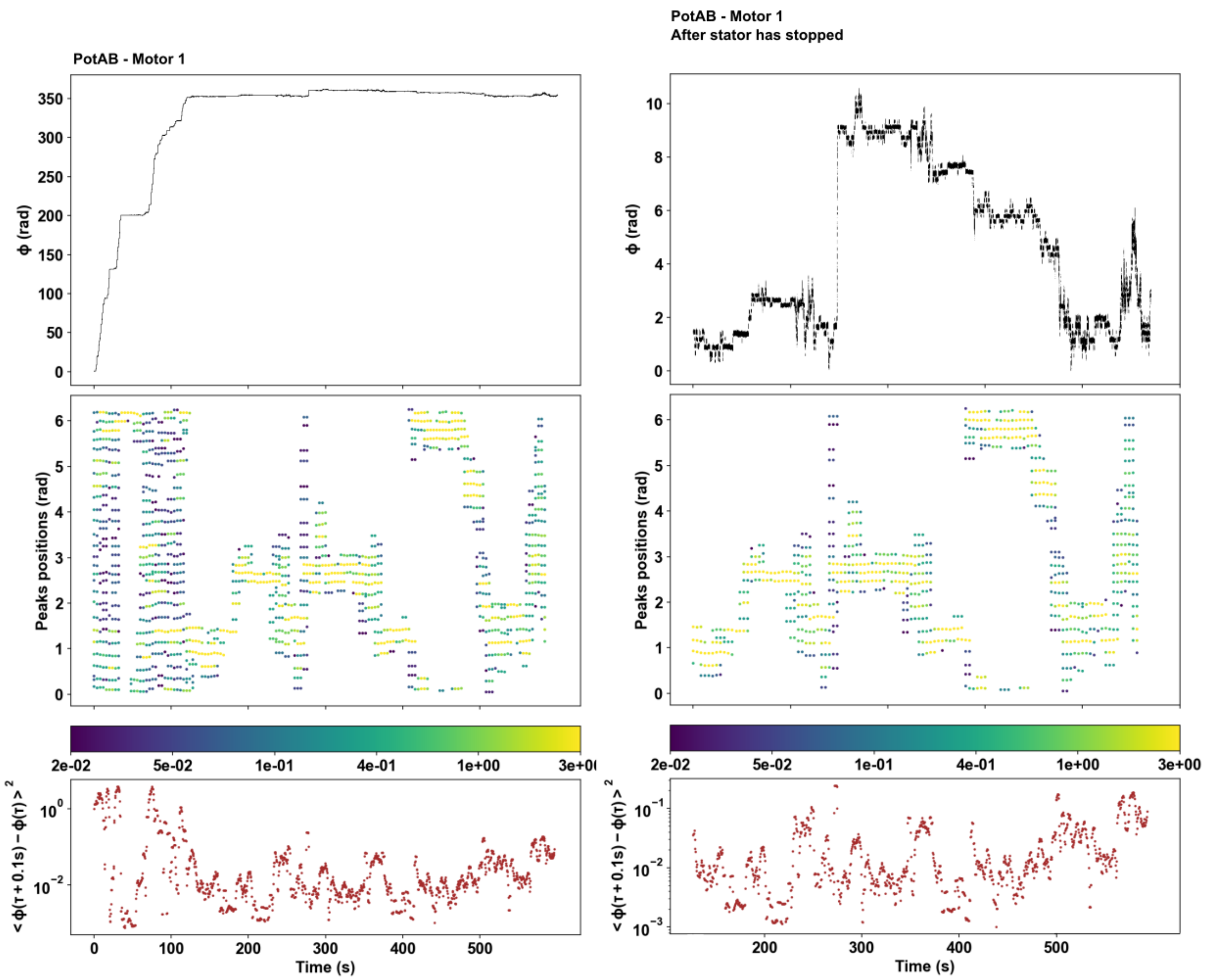


**Supplementary Figure S5**: Same as main Figure 4 but for PotAB Motor 1 (strain MTB24).

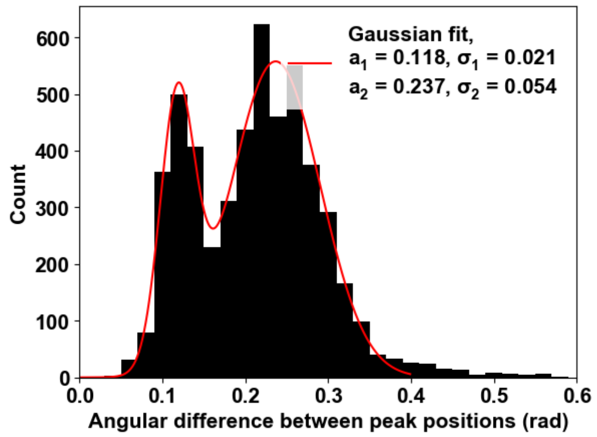


**Supplementary Figure S6:** Histograms over all unpowered cells of the angular differences between adjacent peaks in the histogram of angular positions (represented as dots in Supplementary Figure S5, middle panel). A clear peak appears at 0.237 (~1/26.5 turns) and a second at half that distance.

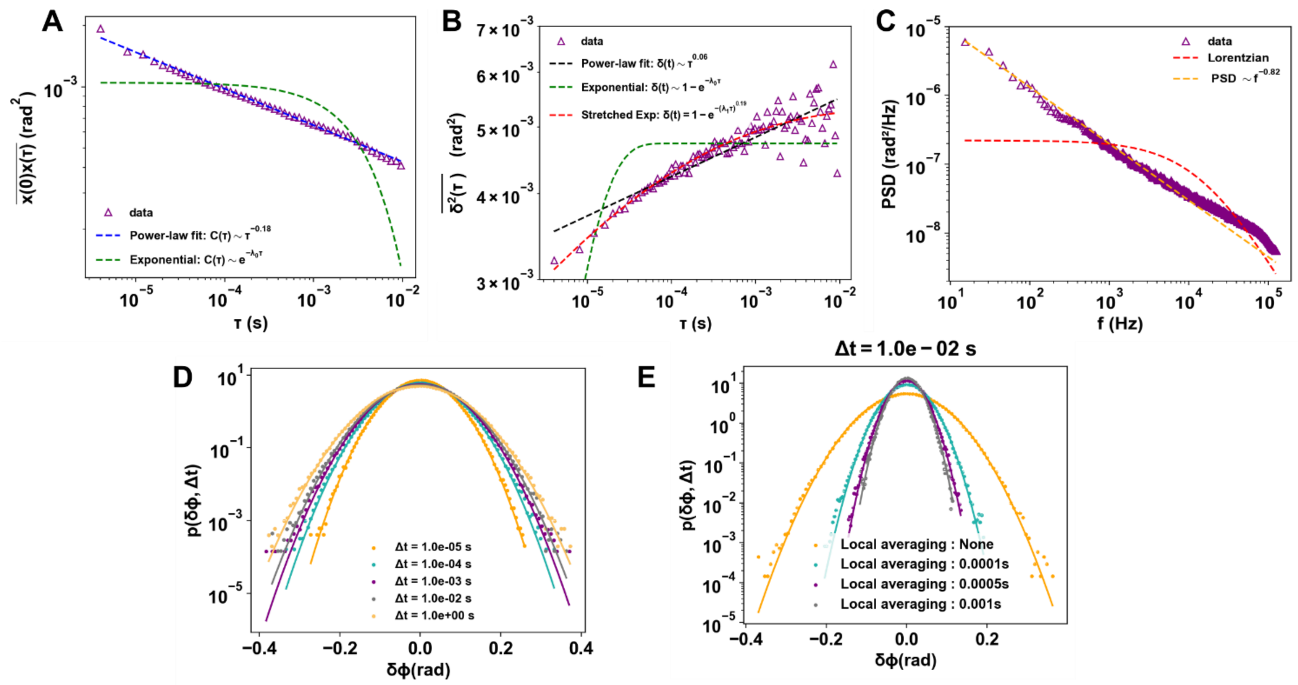


**Supplementary Figure S7:** Detailed view of angular fluctuation from one motor (purple trace in figure 2B). **A.** Time-averaged autocorrelation function of the angular displacement. The data (magenta triangles) cannot be fitted by a single exponential relaxation, but instead follow a power-law decay. **B.** Tie-averaged mean square displacement (TA-MSD) as a function of lag time. A simple exponential relaxation model does not capture the data, whereas a power-law or stretched exponential provides better fits, indicating the absence of a single characteristic relaxation time. A logarithmic MSD would also fit the data. **C.** Power spectral density (PSD) of the angular fluctuations. The spectrum deviates from a Lorentzian shape, further supporting the presence of non-trivial relaxation dynamics. **D.** Probability distributions $p(\delta\phi,\Delta t)$ of the angular displacement $\delta\phi$ at various time intervals $\Delta\tau$ and corresponding Gaussian fits. The distributions remain Gaussian at all timescales. **E.** Probability distributions $p(\delta\phi,0.01s)$ of the angular displacement $\delta\phi$ at a fixed time interval but after pre-processing with various local averaging to remove short-frequency fluctuations. The distributions remain Gaussian.

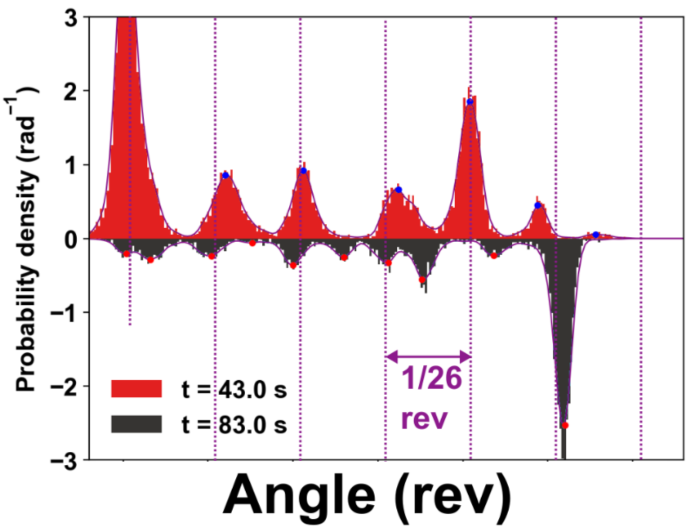


**Supplementary Figure S8**: Enlarged version of the inset of Figure 4A, middle panel. Angular distributions (histogram and Gaussian kernel density estimation) and peaks as measured on 2 non-overlapping windows in ΔMotAB – Motor 4.

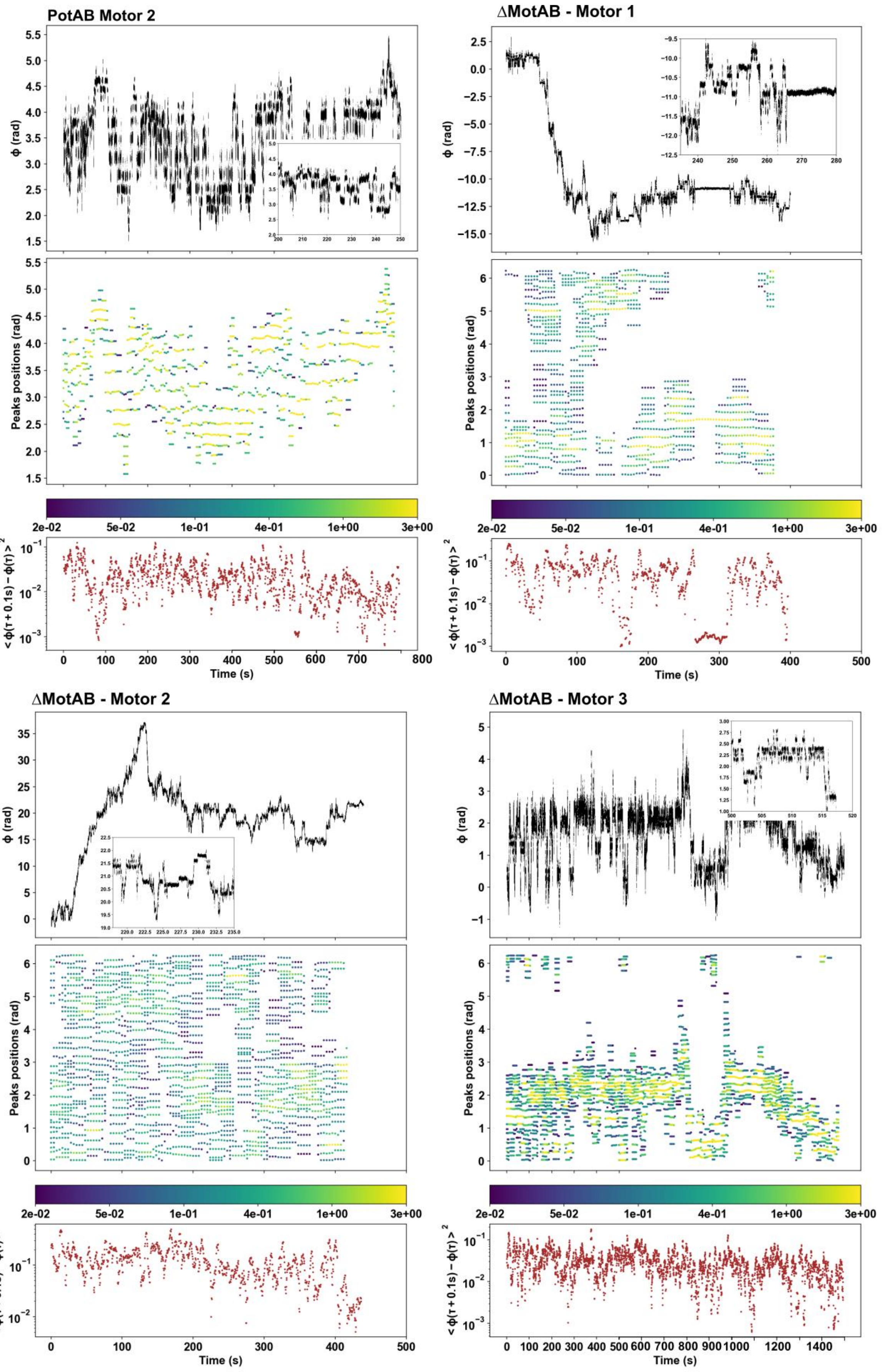


**Supplementary Figure S9**: Same as main figure 4 but for the remaining motors.

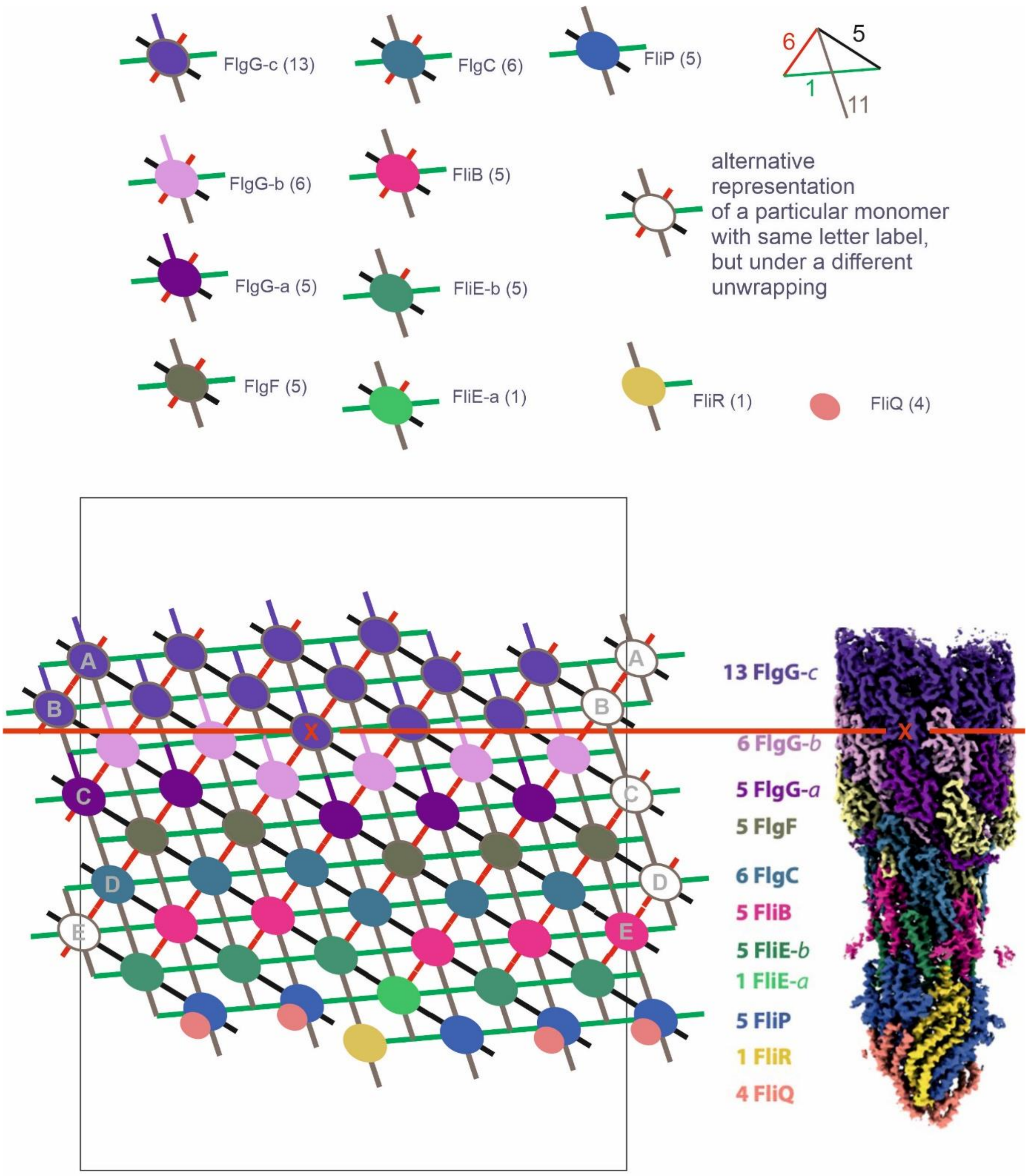


**Supplementary Figure S10:** Representation of the monomer lattice of the flagellar rod, as seen in the structure of [9]. The grey rectangle indicates the rod axis and monomers labelled A-E are duplicated either side of the seam along which the rod has been unrolled. Contacts between monomers on the 1, 5, 6 and 11 start helices are indicated by oblique coloured lines. The horizontal red line indicates the ring where the rod makes close contact with the encircling LP-ring in the flagellar bearing. The lowest FliG-c monomer is marked with a red “x” to illustrate the correspondence between the lattice and the rod structure.

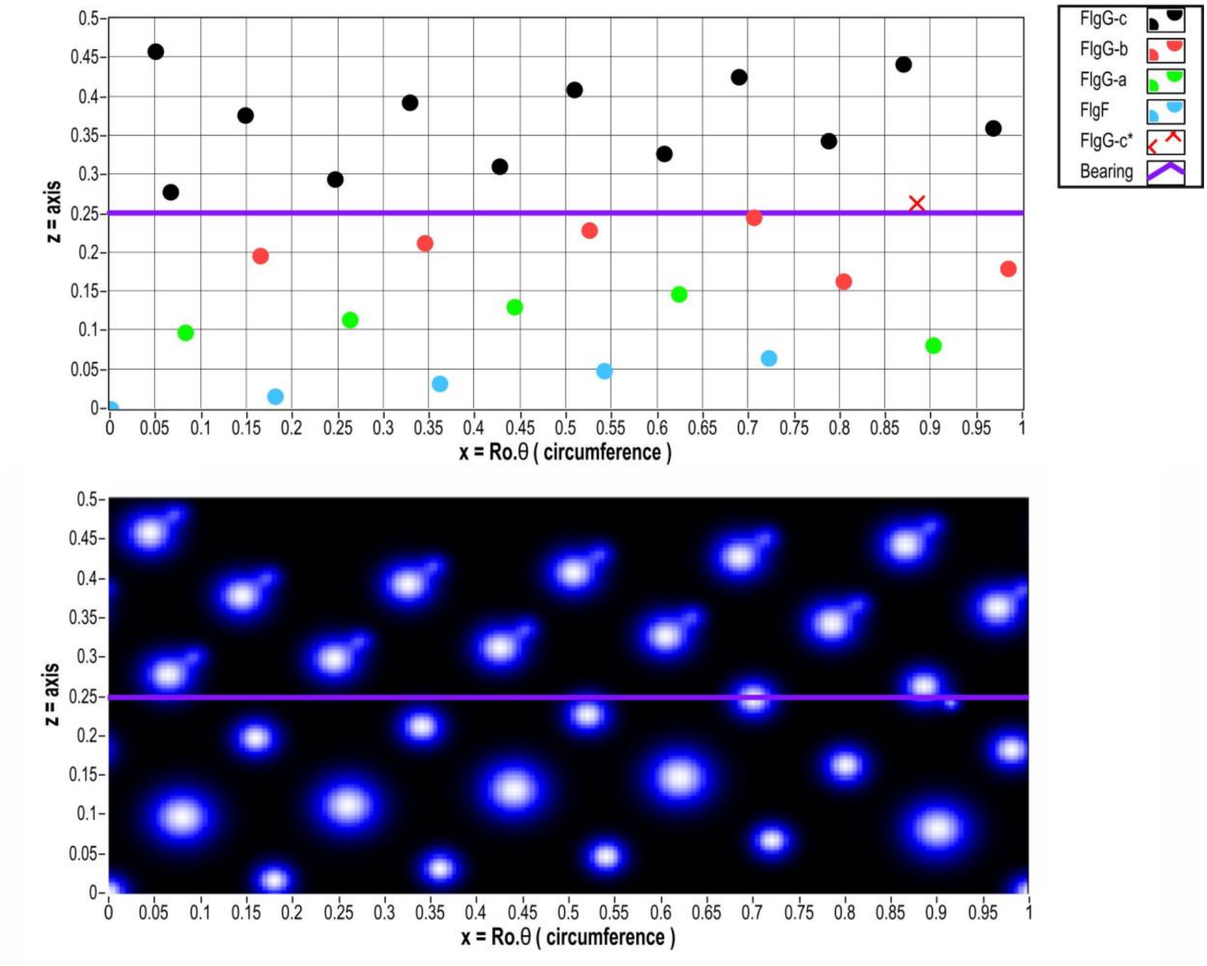


**Supplementary Figure S11:** Model of the monomer lattice of the flagellar rod (top) and a typical interaction potential between the rod and a single LP-ring monomer (bottom). The 4 different distal rod proteins are represented by lattice points of different colour and can have different sets of Gaussian potentials. A unique rod monomer (the first of type FlgG-c), potentially representing a defect or mis-folding, is indicated by a cross in the lattice (top). In this example, the unique monomer is at the axial position of the tight constriction in the LP-ring – the bearing interface (purple horizontal lines) – and has a sharp repulsive feature in its potential (bottom).

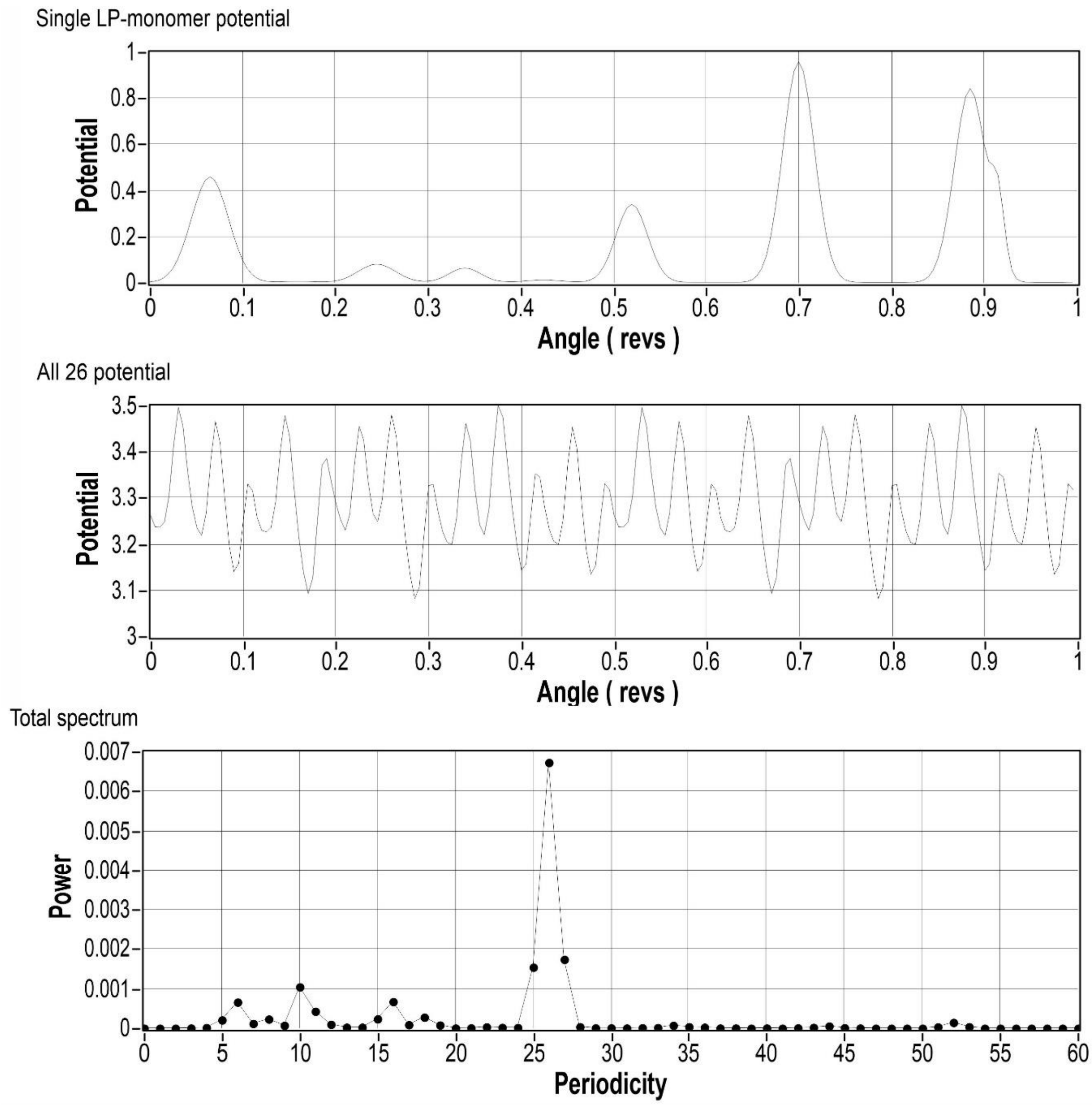


**Supplementary Figure S12:** Single LP-unit potential (top), the total bearing potential (middle) and its Fourier power spectrum (bottom) for the rod potential and LP-ring axial position of Figure S10, in the case where all 26 LP-ring repeating units are identical.   Units of potential and power are arbitrary.

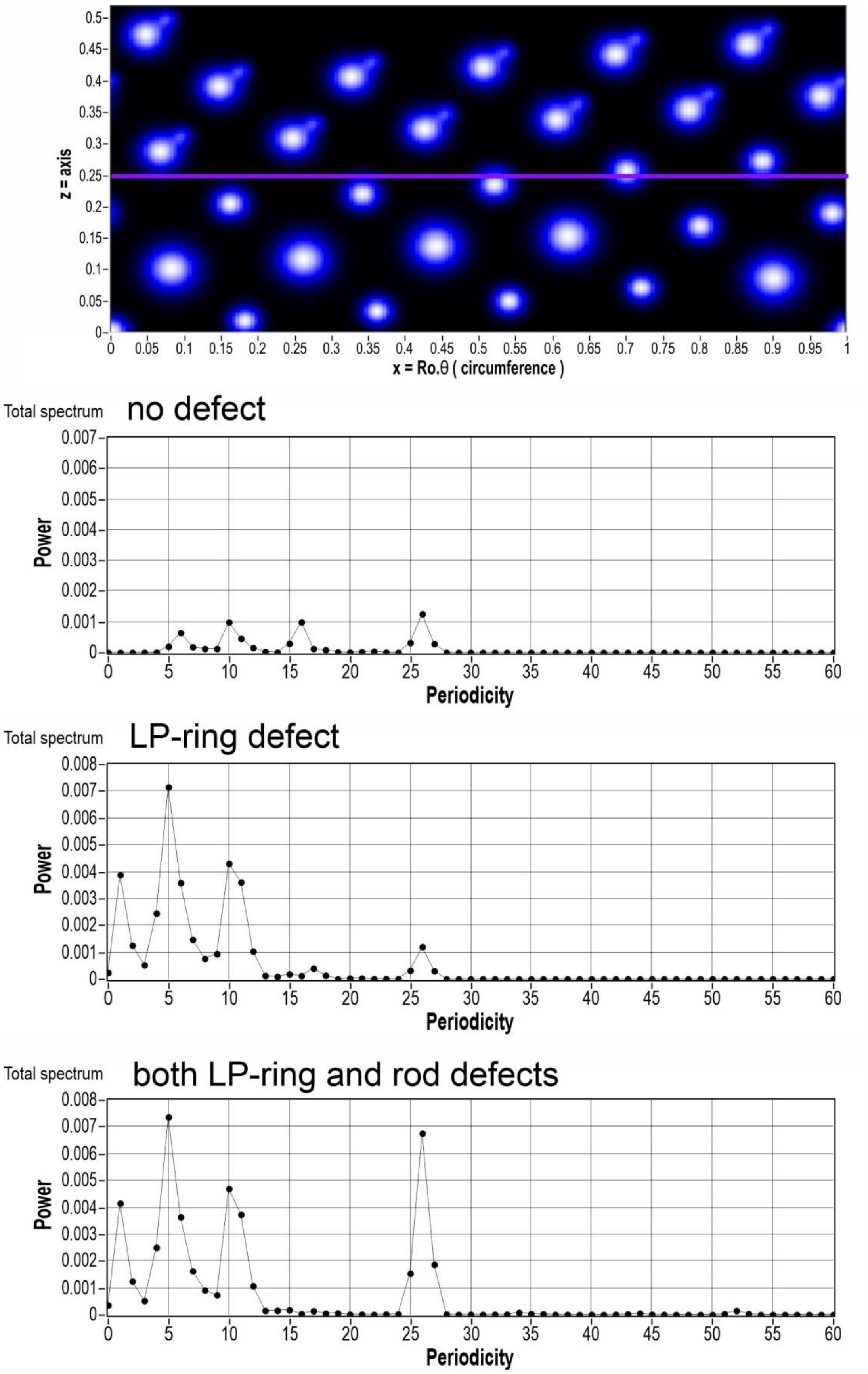
z = axis
x = Ro.θ ( circumference )
Total spectrum
no defect
Power
Periodicity
Total spectrum
LP-ring defect
Power
Periodicity
Total spectrum
both LP-ring and rod defects
Power
Periodicity

**Supplementary Figure S13:** Removing the sharp feature in the rod potential ( in Fig S10, a defect in the unique rod monomer) greatly reduces the 26-fold periodicity (top). Adding a defect in the LP-ring enhances the 5- and 11-fold periodicities (middle, bottom).

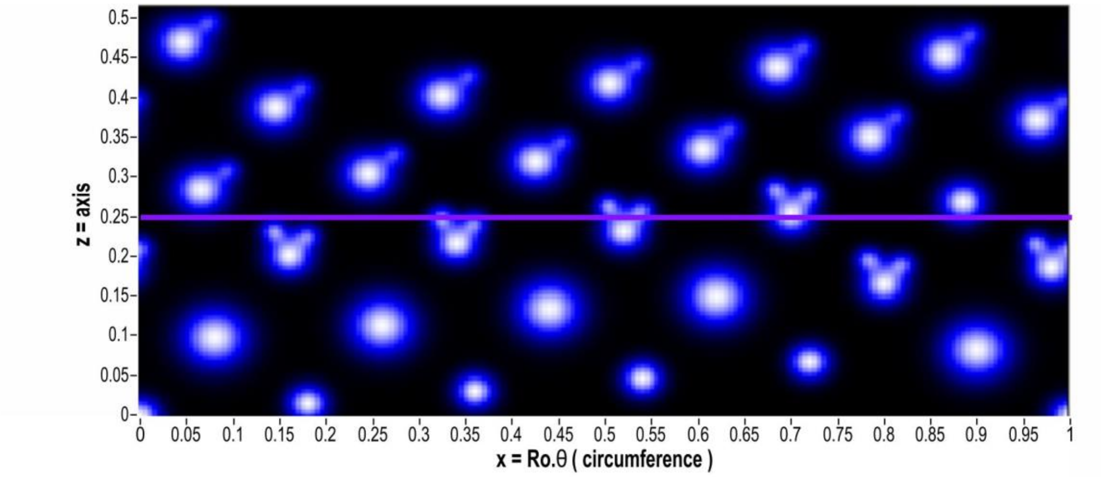


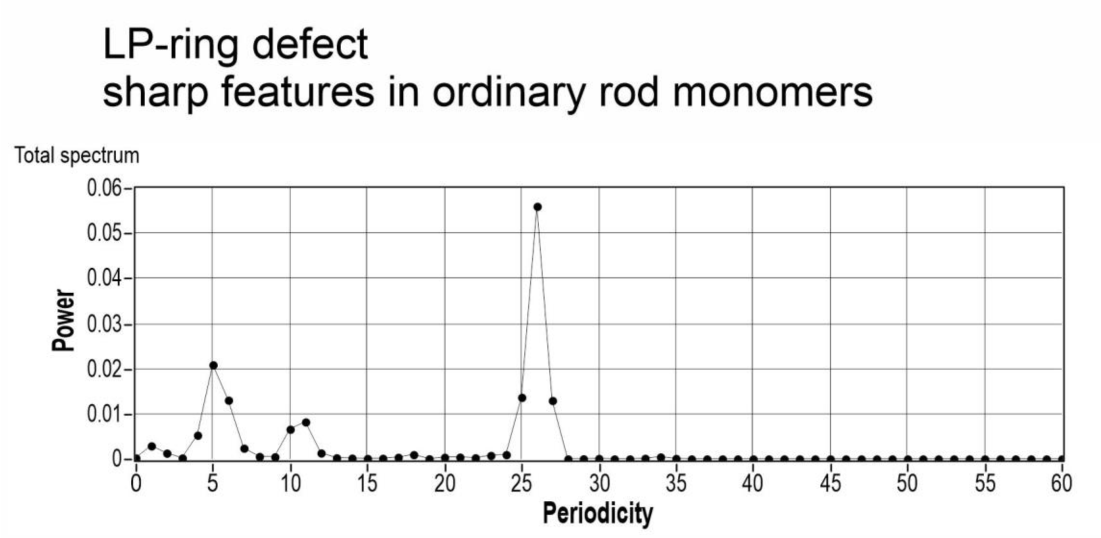


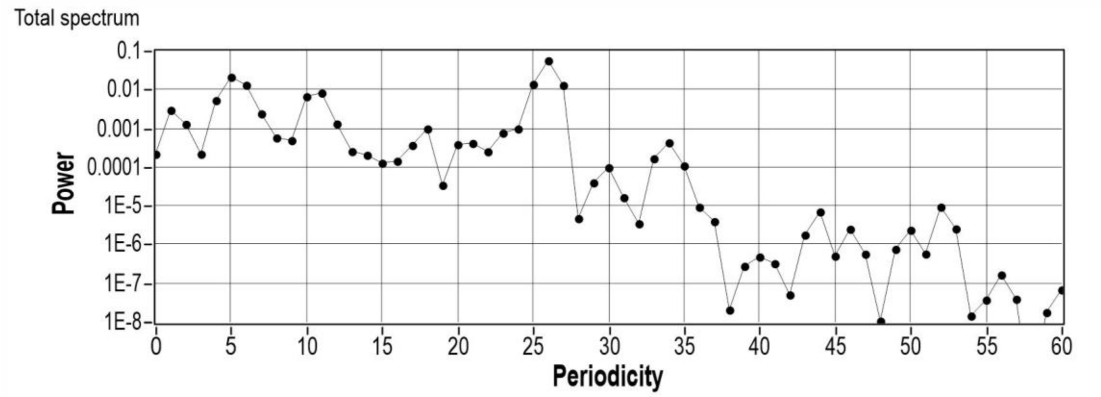


**Supplementary Figure S14:** With a defect present in the LP-ring, sharp features in the rod potential present in all monomers (rather than only defective monomers as in Figs S10-12), also enhance 26-fold periodicity (top). A small peak at 34-fold also emerges in such cases, visible on a logarithmic scale (bottom).
